# Evidence for efficient non-evaporative leaf-to-air heat dissipation in a pine forest under drought conditions

**DOI:** 10.1101/2021.02.01.429145

**Authors:** Jonathan D. Muller, Eyal Rotenberg, Fyodor Tatarinov, Itay Oz, Dan Yakir

**Affiliations:** Department of Earth and Planetary Sciences, Weizmann Institute of Science, 7610001 Rehovot, Israel

## Abstract

- Drier climates predicted for many regions can result in reduced evaporative cooling leading to leaf heat stress and enhanced mortality. To what extent non-evaporative cooling can contribute to plant resilience to the increasingly stressful conditions is poorly known at present.
- Using a novel, high accuracy infrared system for continuous measurements of leaf temperature in mature trees under field conditions, we assessed leaf-to-air temperature differences Δ*T_leaf_*_−*air*_ of pine needles during drought.
- On mid-summer days, Δ*T_leaf_*_−*air*_ remained <1.5 °C, both in trees exposed to summer drought, and in those provided with a supplement irrigation having a 10× higher transpiration rate. The non-evaporative cooling in the drought-exposed trees must be facilitated by low resistance to heat transfer generating large *H*. Δ*T_leaf_*_−*air*_ was weakly related to variations in the radiation load and mean wind speed in the lower part of the canopy, but highly dependent on canopy structure and within-canopy turbulence that enhanced the sensible heat flux *H*.
- Non-evaporative cooling is demonstrated as an effective cooling mechanism in needle-leaf trees, which can be a critical factor in forest resistance to drying climates. The generation of a large *H* at the leaf scale provides a basis for the development of the previously identified canopy-scale ‘convector effect’.

## 1 Introduction

Plants must regulate leaf temperature to control biochemical and physiological processes, such as photosynthesis and water loss, and prevent mortality caused by temperature extremes (Still et al., 2019; Michaletz et al., 2015, 2016; Allen et al., 2010). Leaf temperature, in turn, is a function of leaf traits and environmental conditions. At equilibrium, the energy gained from incoming solar and infrared radiation is balanced by losses due to thermal radiation and latent heat fluxes, and sensible heat exchange (e.g., Gutschick, 2016; Schymanski and Or, 2016). This energy balance can be modified firstly, by leaf traits, such as its size, thickness, structure, and angle, that influence the interception of radiation (Leigh et al., 2012, 2017), and the balance between heat losses through latent and sensible heat fluxes, and secondly by environmental conditions, such as variations in sunlight, temperature, humidity, and wind speed that influence heat transfer. Describing leaf energy balance can be highly accurate under controlled conditions (Gutschick, 2016; Schymanski and Or, 2016; Michaletz et al., 2016), but can be challenging under field conditions. However, identifying the range of variations, and the limits to leaf temperature control resulting from the requirements to maintain energy balance under field conditions is critical to assess vegetation response to climate change, and to scale processes from the leaf to the canopy to ecosystem and larger scales (Still et al., 2021; Bonan, 2008).

In hot and dry ecosystems where plants operate near their limits, maintaining leaf temperature below their thermal threshold, when biochemical and physiological process are damaged, is critical, i.e. 41.5–50.8 °C depending on species and climate zone (O’sullivan et al., 2017; Lancaster and Humphreys, 2020). Indeed, high temperatures can affect a variety of biophysical and biochemical processes (Still et al., 2019; Baldocchi and Penuelas, 2019) like photosynthesis, where they are associated with damage to the functioning of photosystems I & II or the carbon reduction cycle (O’sullivan et al., 2017; Maseyk, 2006; Long et al., 1994; Werner et al., 2002), or an increased vapour pressure deficit and stomatal closure (Smith et al., 2019; Richardson et al., 2020). Not surprisingly, there has been a substantial increase in reports of drought-related tree mortality with rising air temperatures (Allen et al., 2010, 2015).

In water-limited conditions, energy absorbed from incoming radiation that is not used in photosynthesis cannot be readily dissipated through latent heat (transpiration) due to constraints on water availability. Alternatively, this heat can be dissipated through sensible heat. Indeed, semi-arid forests were shown to produce a massive sensible heat flux in spite of having a low surface to air temperature difference by lowering the aerodynamic resistance to heat transfer (*r_H_*) – a property called the ‘canopy convector effect’ that helps dissipate heat from the ecosystem to the atmospheric boundary layer under water-limiting conditions (Rotenberg and Yakir, 2010; Banerjee et al., 2017; Brugger et al., 2018; Kröniger et al., 2018; Zhang et al., 2019). Yet, understanding the evolution of this mechanism at the leaf scale and the regulation of leaf temperature has mostly been discussed in lab experiments (Tibbals et al., 1964; Gates et al., 1965; Michaletz and Johnson, 2006), but not in field conditions.

Leaf cooling through sensible heat is highly dependent on the coupling of leaves and their surrounding air. Indeed, leaf-to-air temperature difference (Δ*T_leaf_*_−*air*_) is often used as a stress indicator in agricultural crops (Kim et al., 2018; Maimaitijiang et al., 2020; Zhang et al., 2019; Song et al., 2017; Long et al., 2006; Jones et al., 2009). A wide range of factors can influence the magnitude of Δ*T_leaf_*_−*air*_ through the heat uptake resulting from the radiation regime and non-radiative heat dissipation which is enhanced through wind penetration into the canopy layer. These factors are affected by the physical location of plants, i.e. elevation and latitude, stand density, or climatic variables such as atmospheric conditions, water supply which affects transpiration rate or large-scale wind patterns (Bonan, 2008; Still et al., 2019; Kim et al., 2016). The interplay between wind and leaf temperature is complex: on the one hand, it can increase leaf water loss because of the transport of moist air away from leaves (Schymanski and Or, 2016; McMahon et al., 2013; Neriah et al., 2014), which increases evaporative cooling. On the other hand, wind can transport sensible heat towards or away from leaves, providing an air cooling mechanism that can decrease water loss, since transpiration is a function of leaf temperature (Schymanski and Or, 2016). Overall, this mechanism is expected to be species-dependent since the reduction in resistance to sensible heat transfer through efficient air flow scales with reduced leaf dimensions (Schymanski and Or, 2016; Taylor, 1975; Geller and Smith, 1982). Indeed, leaf size, geometry and packing of leaves have been shown to be an important factor affecting leaf-to-air heat transfer due to its effect on the leaf boundary layer (Michaletz and Johnson, 2006; Schuepp, 1993; Gates et al., 1965; Tibbals et al., 1964; Schymanski and Or, 2016; Gutschick, 2016). Thin leaves such as those found in conifers have a much lower Δ*T_leaf_*_−*air*_ of 4–8 °C compared to broadleaf species, where Δ*T_leaf_*_−*air*_ reached 10–15 °C (Kim et al., 2018; Leuzinger and Körner, 2007), even in temperate climates where the daytime air temperature rarely exceeds 25 °C (see literature value review in Table S7.1). In agricultural studies, Δ*T_leaf_*_−*air*_ was shown to depend on soil moisture due to the ability of plants to evaporate more water (Siebert et al., 2014). However, since the seminal work of Kim et al. (2016) on ponderosa pine canopies, few studies have followed up on climatic effects on leaf temperature on trees under field conditions, even less so in xeric ecosystems (Lapidot et al., 2019). Furthermore, we are presently unaware of any studies measuring Δ*T_leaf_*_−*air*_ across the height of a canopy, or comparisons between non-water-limited trees that rely on evaporative cooling versus drought-exposed trees that are limited to air-cooling in field conditions.

High precision, direct measurements of leaf temperature under field conditions are a prerequisite to extend our knowledge in this area, but they remain rare (see reviews in Still et al., 2019, 2021). This is particularly so in forests due to the vegetation height and because of the required combined measurement of leaf surface temperatures and that of the surrounding air (Kim et al., 2018). In many studies, thermocouples were used in spite of their limitations (Muller et al., 2021). Therefore, several studies have chosen infrared thermography for leaf temperature (Still et al., 2019; Jones, 2004; Fuchs, 1990). Yet, remaining challenges in using these sensors are the required calibration, knowledge of leaf emissivity (Richardson et al., 2020; Idso et al., 1969) and of background thermal radiation coming from all ecosystem elements (soil, plants, atmosphere). In most studies, emissivity is taken from literature values and background thermal radiation has often either been ignored or simplified using air temperature as a substitute (e.g., Birami et al., 2018), based on the assumption of a small error due to the high emissivity of natural materials (>0.95), an assumption that is not always warranted (Kim et al., 2016). Alternatively, background thermal radiation has been measured using independent sensors (Aubrecht et al., 2016), or empirical correction equations were developed (Kim et al., 2018) using other environmental variables such as air temperature and/or relative humidity. Nevertheless, these methods can lead to substantial measurement errors of up to several degrees, which can be critical in assessing leaf-to-air temperature of similar magnitudes under field conditions. In many studies, air temperature was not measured in the vicinity of leaves, and previous tree canopy-scale studies generally tended to use measurements above the canopy or from nearby meteorological stations, thus amplifying the error of leaf-to-air temperature measurements.

In this study, our objectives were: (a) to measure Δ*T_leaf_*_−*air*_ under field conditions in needle-leaves at a high accuracy using our novel system (Muller et al., 2021); (b) to examine the spatial (across canopy height) and temporal (diurnal) variations of Δ*T_leaf_*_−*air*_ when evaporation is low under drought, or when it is kept high by supplemental irrigation. We hypothesized, that an efficient non-evaporative cooling mechanism at the leaf scale can help maintain low Δ*T_leaf_*_−*air*_, providing resilience to forests undergoing warming and drying climatic trends.

## 2 Materials & Methods

### 2.1 Site description & meteorological measurements

The Yatir forest research site is located in a 2800 ha afforestation of mainly *Pinus halepensis* trees with a height of ~10 m in the dry southern Mediterranean region, at the northern edge of the Negev desert in Israel (31°20′49′′N; 35°3′7′′E; altitude 600–850 m above sea level). Tree density was 300 trees ha^−1^, corresponding to ~50 % crown cover, and the lowest branches were at ~2 m agl. Mean annual global radiation was 238 W m^−2^, while average air temperatures for January and July are 10 and 25.8 °C respectively, with a mean annual potential ET of 1600 mm, and mean annual precipitation of 285 mm (Rotenberg and Yakir, 2010; Tatarinov et al., 2016; Qubaja et al., 2019). Measurements were done during the extended dry season (June to October) typical for the semi-arid southern Mediterranean climate zone.

The research site contains an eddy covariance flux tower operating since 2000, whose above-canopy environmental sensors (15 m agl; air temperature (°C), RH (%), horizontal wind speed component at 18.7 m agl (m s^−1^); R3-100 3D sonic anemometer from Gill Instruments, Lymington, United Kingdom), shortwave (0–4000 nm (W m^−2^); CM21, Kipp & Zonen B.V., Delft, The Netherlands) and longwave radiation (4–50 μm (W m^−2^); Eppley, Newport RI) were used for auxiliary measurements of meteorological conditions during our consecutive measurement periods (Table 1) on a half-hourly timescale.

**Table 1.**
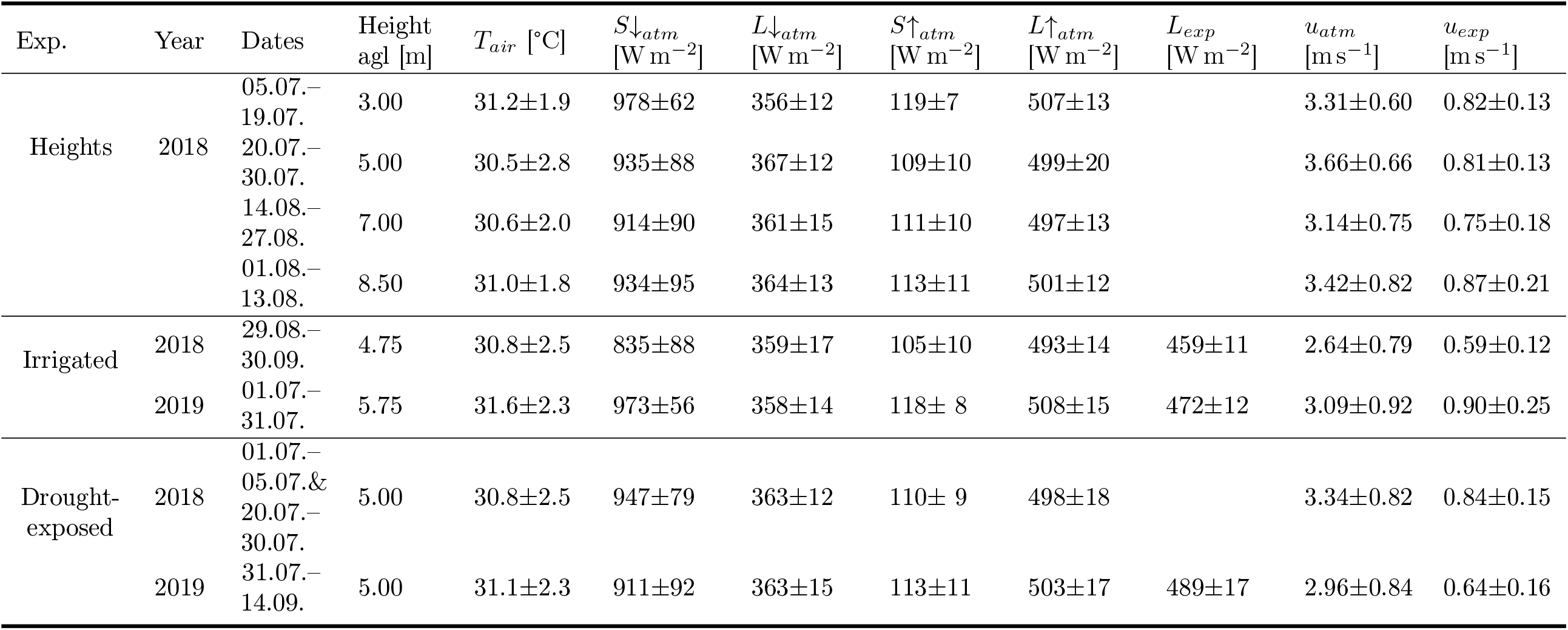
Mean and standard deviations (in parentheses) of midday conditions (10:00-14:00) during consecutive measurements of leaf temperatures in summers 2018 & 2019 with standard deviations, showing conditions in the experimental height (‘exp’) and in the atmosphere, measured above canopy at 15m agl (‘atm’). Experiments consisted of measurements in different heights in 2018, and in an irrigated and drought-exposed plot in 2018 & 2019. Columns show the experiment (Exp), year and exact dates of measurements, height above ground level, air temperature *T_air_* [°C], down- (↓) and up-welling (↑) solar (*S*) and longwave thermal (*L*) radiation above canopy [W m^−2^] and *L_atm_* [W m^−2^], respectively, longwave background radiation in the experimental height *L_exp_* [W m^−2^], wind speed above canopy and in the experimental height *u_atm_* [m s^−1^] and *u_exp_* [m s^−1^], respectively.

Midday conditions (10:00-14:00) were similar during the consecutive measurement periods: air temperatures were ~31 °C, above-canopy incoming short- and longwave radiation were ~929 and 362 W m^−2^, respectively, while above-canopy wind speed averaged 3.22 m s^−1^ and VPD 2.72 kPa. These values, summarized in Table 1, show that conditions during summer drought remain stable and comparable, even though measurements were not taken simultaneously (i.e. the period in 2018 & 2019 was not identical).

Finally, measurements from an experimental plot active since the spring of 2016 were available, where gas exchange chambers were used to measure assimilation and transpiration of multiple leaves on 4 twigs per chamber in ~5 m agl in cycles of 1h under summer supplemental irrigation and drought-exposed (i.e., control) conditions (Preisler, 2019). Our system was deployed to measure *T_leaf_* on neighbouring twigs of the ‘irrigated’ and ‘drought-exposed’ experiments (Table 1), and those simultaneous measurements were used for data analysis.

### 2.2 Leaf and air temperature: Measurement principles

Thermal infrared cameras record thermal radiation from different sources: first, that emitted by the object of interest (*L_obj_*) with emissivity *ε_obj_*. Second, the radiation of the surrounding background (*L_bg_*) that the object reflects according to 1 − *ε_obj_* (with both object and reflected radiation attenuated through the air column by a transmissivity coefficient *τ*). Third, the thermal radiation of the air column (*L_air_*) between the sensor and the object emitting according to 1 − *τ*, which is affected by the air temperature, humidity and aerosols content (Fig. 1). Hence, the total measured longwave radiation (*L_tot_*) can be described by Eq. 1 (Aubrecht et al., 2016; Incropera et al., 2006; FLIR, 2011), and *τ* is assumed to be 1 when objects are close to the infrared camera:

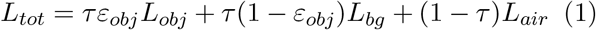

**Fig. 1.**
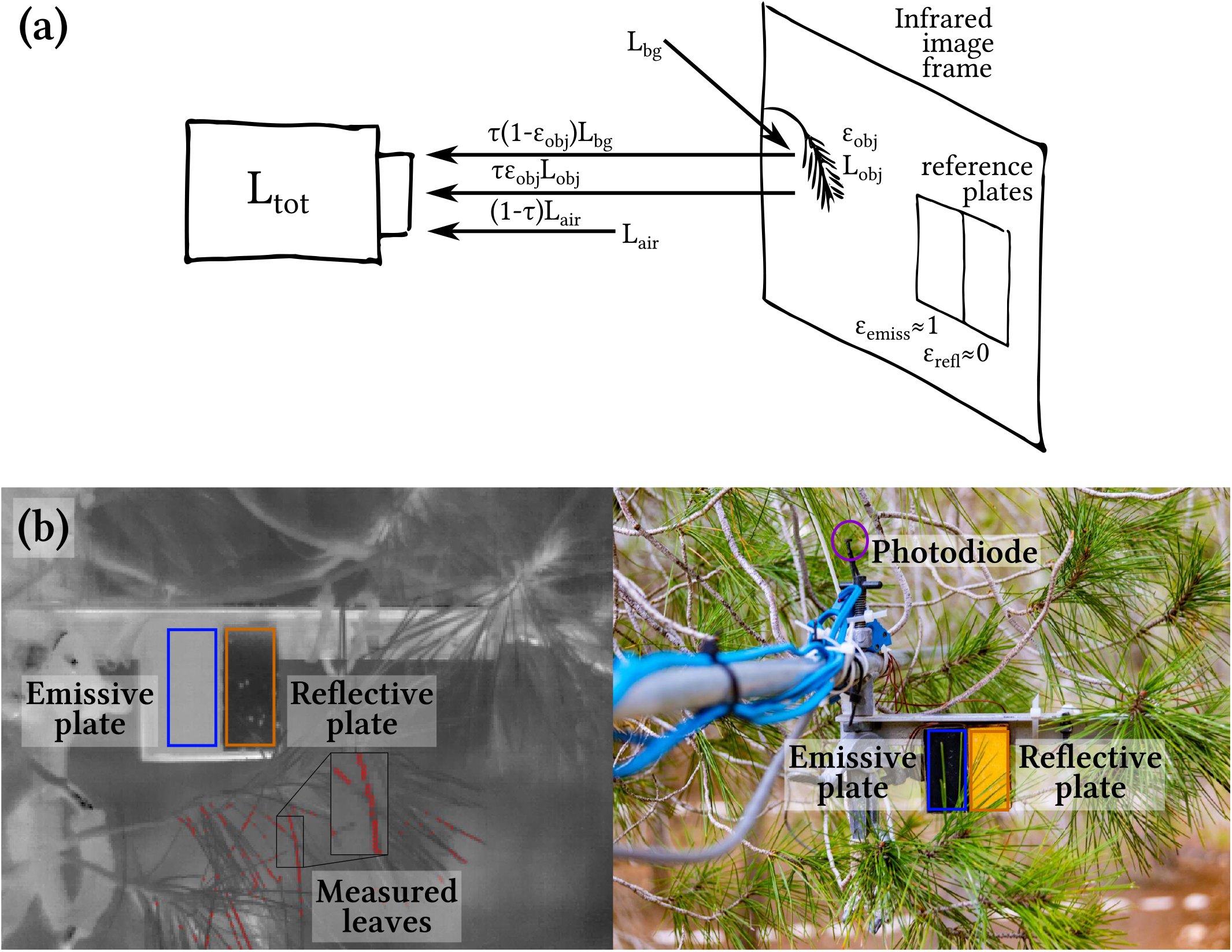
(a) Illustration of the total radiation reaching an infrared camera (*L_tot_*) and the relevant terms it receives, including a set of two reference plates with low (*ε_refl_*) and high (*ε_emiss_*) emissivities, respectively, in its field of view. Part of *L_tot_* is emitted by the object (*L_obj_*), by the air column (*L_air_*) between it and the camera which attenuates it (*τ*), and part is a reflection of the background (*L_bg_*) coming from surrounding objects and the sky. (b) Left side: Example infrared image where dark shades are colder, where red points (cf. zoomed-in square) show examples of pixels identified as leaves by automatic scripts; Right side: Setup of the infrared camera’s picture frame. Polygons show the reference plates used for camera offset correction (emissive plate, blue) and background thermal radiation (reflective plate, orange) and a photodiode (purple) measured the incoming shortwave radiation.

Infrared cameras calculate the apparent object temperature (*T_ap_*) using operator-provided fixed values for *ε_obj_*, *L_bg_*, *L_air_* and *τ*. When the objects (or parts of them) in an infrared image have different emissivities, the temperature of those pixels has to be adjusted. It is common practice to set the camera settings of *ε* and *τ* to 1.00 to simplify corrections in post-processing. Systematic camera error can be corrected by correlating the infrared measured temperature of a reference plate with a high emissivity (*ε_emiss_* ≈ 1; Fig. 1) with its independently measured temperature (Kim et al., 2018).

In ecological applications, the background thermal radiation *L_bg_* has often been ignored due to the high emissivity of natural materials and relatively small contribution of *L_bg_*. This is sufficiently accurate where only relative temperatures are required, but can lead to substantial errors in determining the actual temperature (Kim et al., 2016). In the present study, we employed a low emissivity plate (*ε_refl_* ≈ 0, Fig. 1) with an integrated thermocouple, placed in the image frame, to measure *L_bg_*. The purpose of the embedded thermocouple is to account for the residual thermal radiation from the reflective plate observed by the thermal camera, as its emissivity is not exactly zero (Table 2). We were therefore able to measure *L_bg_* near the sampled object in the same spectral range as the thermal measurements according to Eq. 2:

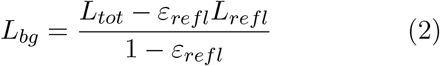

where *L_tot_* is calculated using the Stefan-Boltzmann law 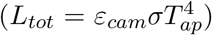 from the apparent temperature (*T_ap_*) of the reflective plate reported by the infrared camera, using the Stefan-Boltzmann constant *σ* and the camera emissivity setting *ε_cam_* (set to 1). *ε_refl_* is the emissivity of the highly reflective plate (Table 2) and the thermal radiation emitted by it is 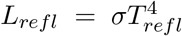, where *T_refl_* is the independently measured temperature of the reflective reference plate (using a calibrated thermocouple). Finally, the corrected object temperature can be calculated by applying the resulting *L_bg_* in Eq. 1 (Muller et al., 2021).

**Table 2.** Mean hemispherical directional emissivity with number of samples (N) and standard deviation of multiple samples of different materials measured in the lab using our custom-made system (Vishnevetsky et al., 2019). For an emissivity *ε* > 0.90, the uncertainty is ≤ 0.5% of the measured *ε*, for *ε* < 0.95, the uncertainty is ≤ 0.4%, for *ε* ≤ 0.1, the uncertainty is ~28.6%

| Sample Materials | N | Emissivity |
| --- | --- | --- |
| Branch bark | 4 | $0.896 \pm 0.014$ |
| Emissive coating; <2018-09-06 | 1 | 0.984 |
| Emissive coating; >2018-09-06 | 1 | 0.977 |
| Reflective coating | 1 | 0.023 |
| Pine needles (dry) | 3 | $0.933 \pm 0.010$ |
| Pine needles (fresh; 1-3 yrs old) | 4 | $0.947 \pm 0.003$ |
| Topsoil (Yatir research site) | 4 | $0.933 \pm 0.006$ |

### 2.3 Leaf and air temperature: Field measurements

Leaf temperatures were measured during the peak of summer drought, consecutively in different heights within the canopy in summer 2018 (3, 5, 7 & 8.5 m); as well as in ~5 m (middle of the canopy) in a drought-exposed and experimentally irrigated plot in summers 2018 & 2019 (Measurement periods & location, cf. Table 1, map: Fig. S4.1). The mast and camera were installed and raised manually from the ground. Once extended, it was oriented so that the reference plates and photodiode would be near the leaves that were to be measured. As an unavoidable consequence, the branches in different heights were not all exactly above each other, resulting in some differences in radiative exposure of the measured leaves, as could be expected under the in-situ field conditions. In the following text, leaf temperature will refer to the mean temperature of multiple needles on the same twig (as seen in Fig. 1b).

Thermal infrared measurements of *T_leaf_*, *T_emiss_*, *T_refl_* and *L_bg_* were done using an infrared camera (7.5–13 μm; FLIR A320; FLIR Systems, Wilsonville, Oregon, United States) facing north installed on a mast, triggered automatically every 5 min (every 15 min before 2018-08-01) using a camera setting of *ε* = 1.00 and *τ* = 1.00 due to the close distance of <65 cm between the infrared camera and the leaves (setup cf. Fig. 1). Other camera parameters are dropped due to *ε* = 1 and *τ* = 1. At this distance, pixels were ~0.4 mm wide using a 15° lens, compared to a diameter of needle-leaves of ~0.8–1 mm, which allowed us to avoid mixed values between the background (sky, branches) and the leaves (leaves). The final temporal resolution was lower than 5 min because the camera had to be restarted every few hours to circumvent crashes of the internal camera software. Measurements were averaged to half hours, i.e. equivalent to the typical time step for eddy covariance flux data. An arm extending from the infrared camera held branches in a fixed position to minimise movement from wind and simplify image analysis. Systematic camera error, i.e. the offset from the real surface temperature, was corrected using measurements of a reference aluminium plate with a highly emissive coating (*Metal Velvet™* coating; Acktar Ltd., Kiryat-Gat, Israel). For measurements of *L_bg_*, we used a reference plate with a high reflectivity (Infragold coating; Labsphere Inc., North Sutton NH, USA; installed on 2018-09-06) alongside it. Both plates had a thickness of 3 mm, and each contained a thin thermocouple installed in the middle of the plates, 0.5 mm from the coated surfaces in a tiny drilled hole filled with thermal grease. The plates were insulated by a piece of wood 2 cm thick on the back to prevent undesired heating from the air. Their hemispherical directional emissivities (at a right angle to the plates), as well as that of natural materials present in Yatir forest (needle-leaves, branches, soil) were measured in the lab using our special emissivity measurement instrument (Vishnevetsky et al., 2019) at an uncertainty of <0.5 % for an emissivity >0.90 (Table 2). Note that only mean emissivity values for fresh needles are indicated, since no differences due to leaf age or leaf water content could be identified (i.e. difference <0.005). The total temperature measurement error of our system was ±0.25 °C (Muller et al., 2021), as calculated from the combined errors of each sensor using the log derivative method (Fritschen and Gay, 2012). This is much more accurate than the previously reported IR camera accuracy of ±2 °C (FLIR, 2011).

A photodiode sensitive in the PAR range (400–700 nm; G1118, Hamamatsu Photonics K.K., Japan), calibrated against a commercial PAR sensor (PQS1, Kipp & Zonen, Delft, The Netherlands) and a short-wave full-range pyranometer with a 0–4000 nm range (CM21, Kipp & Zonen B.V., Delft, The Netherlands), was employed to measure incoming solar radiation near the leaves at 1 Hz. The light reaching this photodiode was assumed to be the same as in the entire field of view of the thermal camera.

At a distance of ~2 m from the measured leaves, a set of two 3D sonic anemometers (Windmaster Pro, Gill Instruments, Lymington, UK) were installed vertically (1.5 m distance from each other) on a secondary mast, where the middle point between them was at the same height as the measured leaves. Each sonic measured three components (*u*, *v* & *w*), from which the horizontal wind speed component (normalised to the direction and represented as *u* from here on) and shear velocity (*u*_∗_) were calculated at 6 min time steps and averaged to ½h time steps (using EddyPro 7.0.6; LI-COR, Lincoln NE, USA). Then, the means of *u* and *u*_∗_ between the two anemometers were used for further analyses. Data was logged on a field computer using a LabVIEW (National Instruments; Austin, Texas USA) software. For precise air temperature measurements, a calibrated fine T-type thermocouples were installed in Young radiation shields (Model 41003; R. M. Young Company, Traverse City MI, USA) in the same height and on the same mast as the anemometers. Calibration of all thermocouples was done in stirred water of a known uniform temperature to remove differences of individual thermocouple junctions, as well as against a precise air temperature thermistor in a radiation shield. The accuracy and precision of the thermo-couples was 0.1 °C, that of the FLIR A320 infrared camera was ±1 °C and 50 mK at 30 °C, respectively (FLIR, 2011).

### 2.4 Data extraction & outlier removal

The high-resolution raw infrared temperature data contained in the radiometric image files captured by FLIR cameras was extracted using a custom script (Muller and Segev, 2020). Needle-leaves were then identified using local temperature minima of cold leaves compared to their blurred warmer background (Muller et al., 2021). This method was compared to manual sampling to assess its robustness (Section S1). The median and standard deviations of all the detected pixels in each category (i.e. needle-leaves and reference plates) were calculated for each infrared image. Note that this provided a median temperature of needle-leaves on a twig, denoted as ‘leaf’ temperature.

Thermal sensors suffer from systematic sensor drift over time, and the usage of a reference plate with a high emissivity is recommended to account for this (Kim et al., 2018). This correction was performed using night-time data only (*R*^2^ = 0.99, *P* < .001, *RMSE* = 0.13°C) in order to avoid the potential inaccuracy caused by temperature differences between the reference plates’ surfaces and the built-in thermocouples caused by fast fluctuations of sunlight on the plates (Muller et al., 2021). Outliers resulting from sensor issues related to crashes of the infrared camera’s internal software, thermocouples, dust or humidity on the reference plates leading to > ±2 °C differences between both reference plate thermocouples were removed (<1 % of data). Before the installation of the reflective plate on 2018-09-06, only the emissive plate was used to calibrate the IR camera readings of the leaf surface temperatures using a linear correlation between the built-in thermocouple and the IR reading.

## 3 Results

### No effect of transpiration rates

Δ*T_leaf_*_−*air*_, measured using the high precision temperature measurement system, and transpiration (TR), measured using automated branch-scale gas exchange chambers, during the 2018-2019 study period are summarized in Figure 2. At SWR of 800–900 W m^−2^, TR reached maximum values of 0.12 ± 0.07 mmol m^−2^ s^−1^ in the drought-exposed plot. This translates to a latent heat (LE) of ~5 ± 3 W m^−2^ under the highest SWR and net absorbed radiation levels (*R_n_* of 398 ± 8 W m^−2^). TR reached >10× higher values of 2.34 ± 0.67 mmol m^−2^ s^−1^ in the irrigated plots, i.e. LE ~103 ± 29 W m^−2^ under a similar *R_n_* of 387 ± 17 W m^−2^). Δ*T_leaf_*_−*air*_ values showed a clear dependence on incoming solar radiation SWR (drought-exposed: Δ*T_leaf_*_−*air*_ = 2.89 *exp*(−0.006 *SWR*^0.84^), *R*^2^ = 0.81; irrigated: Δ*T_leaf_*_−*air*_ = 3.40 *exp*(0.001 *SWR*^1.18^), *R*^2^ = 0.88), showing an approach to saturation above 600 W m^−2^ (Fig. 2a; see also Fig. S3.1).

**Fig. 2.**
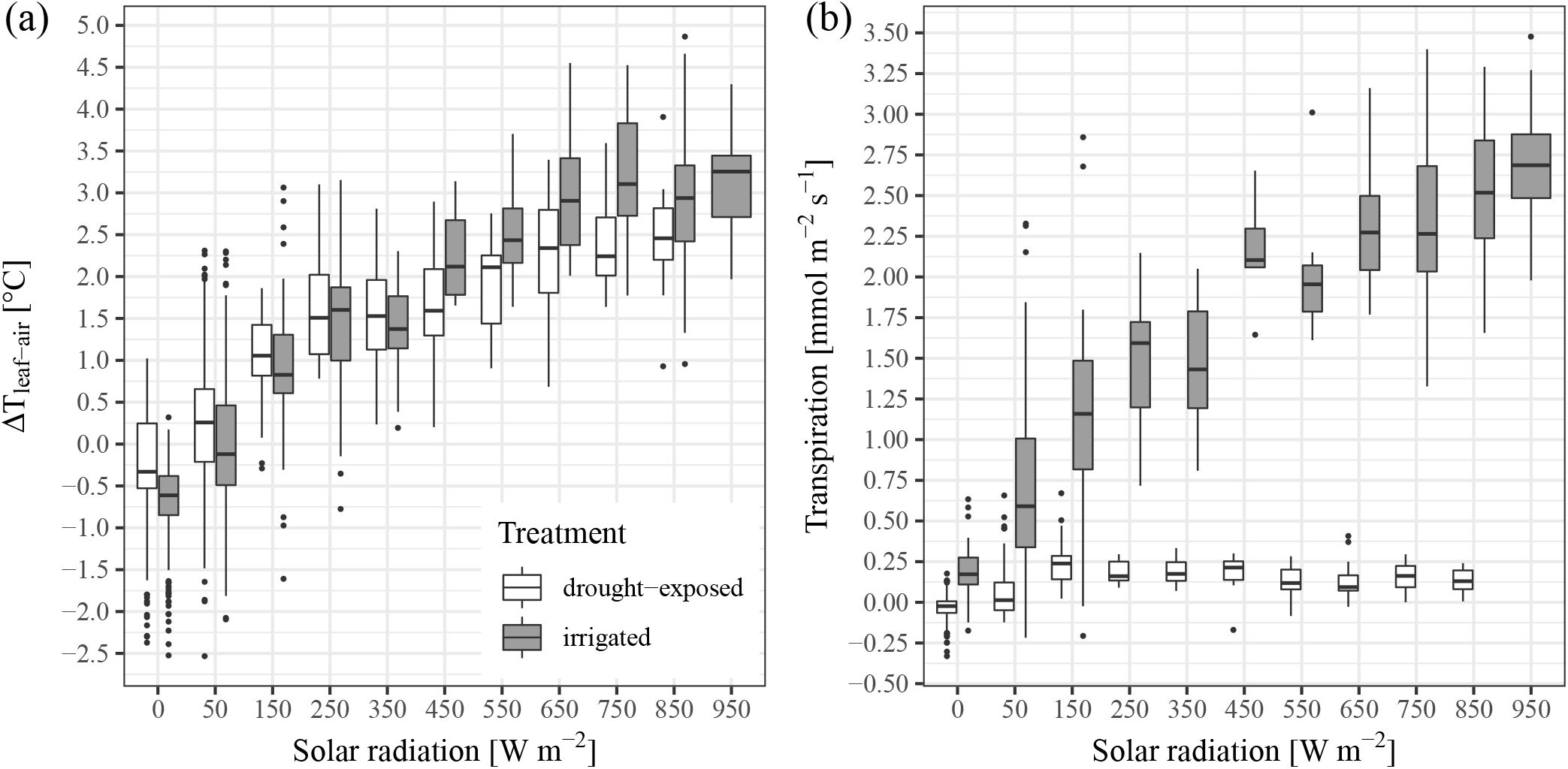
Relationship between leaf-to-air temperature difference Δ*T_leaf_*_−*air*_ and (a) shortwave radiation, with significance of differences between treatments, and (b) transpiration in bins of SWR of 100 W m^−2^, measured in summers 2018 & 2019 in irrigated (yellow) and drought-exposed trees (black) in 5 m agl. Bars represent standard deviations for each bin, asterisks show significance of differences between experimental treatments.

The mean canopy-layer air and leaf temperatures (~5 m agl) as well as Δ*T_leaf_*_−*air*_ are reported in Table 3. The results show that in spite of similar *R_n_* and large differences in TR, Δ*T_leaf_*_−*air*_ was similar on average in both plots during the day. We are particularly interested in Δ*T_leaf_*_−*air*_ under high radiation load (SWR >600 W m^−2^), and as shown in Fig. 2, mean Δ*T_leaf_*_−*air*_ reached values of 2.4 ± 0.6 and 3.0 ± 0.8 °C (*P* < .001), for drought-exposed and irrigated trees, respectively. In contrast to expectations, Δ*T_leaf_*_−*air*_ was even slightly higher in the high transpiration irrigated plot, associated with slightly lower air temperature, which is likely due to the wetter soil below it (Table 3).

**Table 3.** Mean of daytime maxima (10:00-17:00) and night-time minima (21:00-05:00) of *T_leaf_*, *T_air_* and *T_leaf_*_−*air*_, in [°C]

| Value | Time | Drought-exposed | Irrigated | P |
| --- | --- | --- | --- | --- |
| $T_{leaf}$ | Day | $34.1 \pm 1.8$ | $32.9 \pm 2.1$ | 0.02 |
| | Night | $18.4 \pm 1.9$ | $17.8 \pm 2.1$ | 0.16 |
| $T_{air}$ | Day | $32.9 \pm 2.1$ | $32.0 \pm 2.1$ | $<.001$ |
| | Night | $18.2 \pm 1.5$ | $17.7 \pm 1.9$ | $<.001$ |
| $\Delta T_{l-a}$ | Day | $2.5 \pm 0.7$ | $2.5 \pm 1.4$ | 0.96 |
| | Night | $-0.7 \pm 0.8$ | $-1.2 \pm 0.6$ | $<.001$ |

A significant relationship between TR and Δ*T_leaf_*_−*air*_ was exhibited in irrigated trees (Fig. 2b; Δ*T_leaf_*_−*air*_ = −0.97 + 1.61*TR*, *R*^2^ = 0.96, *P* < .001), but not in drought-exposed trees where Δ*T_leaf_*_−*air*_ increased to similar levels, but with a near lack of transpiration (Figure 2b; *R*^2^ = 0.01, *P* = 0.78). The increase of Δ*T_leaf-air_* seems to be largely driven by SWR, which also drives the increases in TR in irrigated trees. Therefore, the remarkably similar maximum Δ*T_leaf-air_* value was observed in the two treatments in spite of the ~10× difference in TR values. This indicated that evaporative cooling in itself does not prevent leaf heating (i.e., Δ*T_leaf_*_−*air*_ increased in spite of large increase in TR; Fig. 2b).

### Factors affecting leaf to air temperature differences

Δ*T_leaf_*_−*air*_ and the factors that can influence it were examined in more detail in the drought-exposed, i.e. control, plot and are summarized in Fig. 3. Shortwave radiation measurements (SWR) highlight the differences in shading in the different heights (Fig. 3c), which was associated with the orientation of the mast and the complex canopy structure in this field study. Therefore, the infrared camera measured more exposed leaves of the exposed outer parts of the canopy (‘canopy skin’) at 7 m, where SWR levels were similar to the ones above canopy (up to ~1000 W m^−2^). At 3, 5 & 8.5 m, canopy shading reduced mean SWR input to a maximum ½h mean of 550 W m^−2^.

**Fig. 3.**
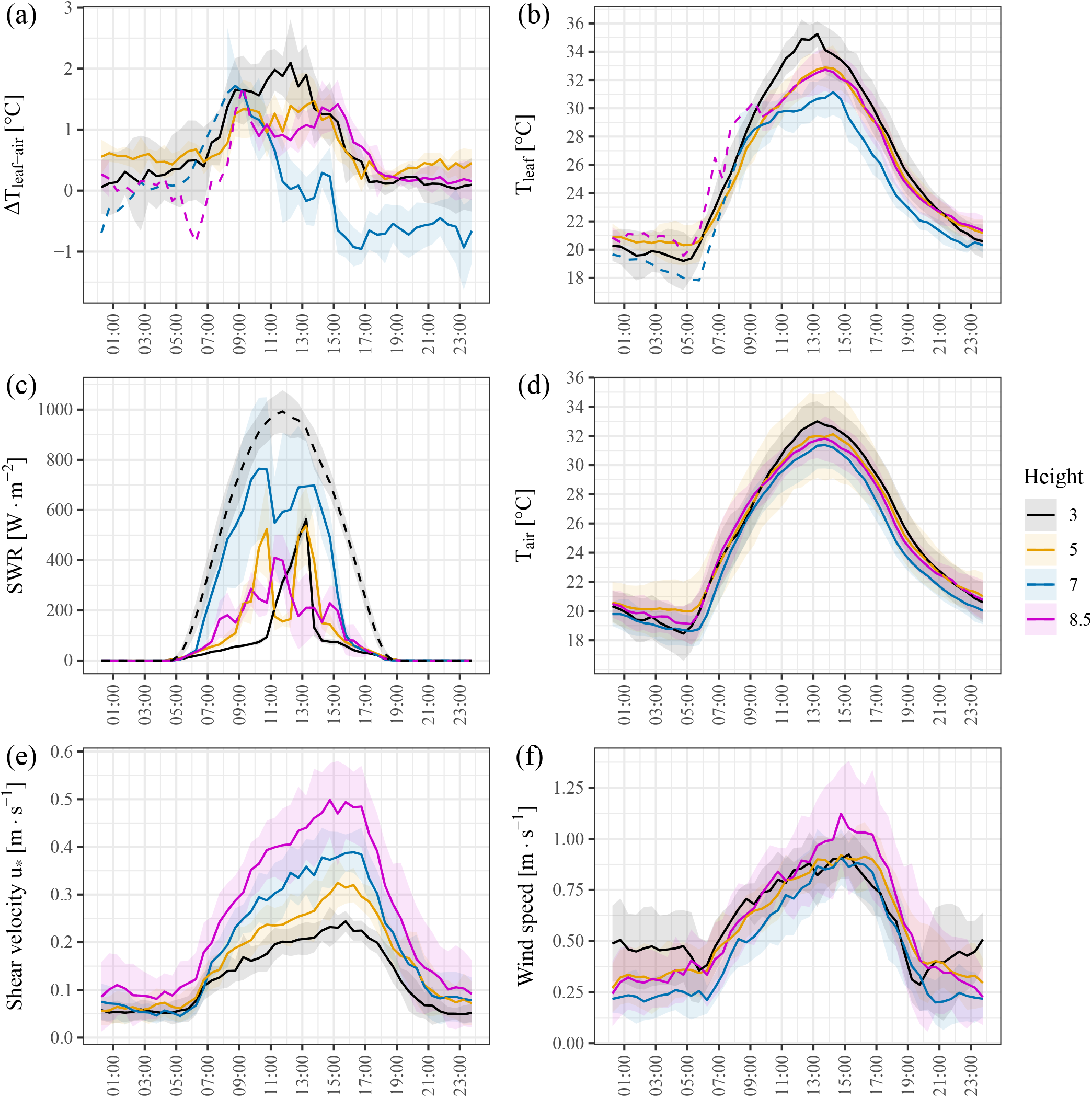
Diurnals of half-hourly means over 2 weeks in the drought-exposed plot of (a) leaf-to-air temperature difference Δ*T_leaf_*_−*air*_ [°C], (b) leaf temperature *T_leaf_* [°C], (c) incoming shortwave radiation SWR [W m^−2^] with above-canopy SWR (dashed black), (d) air temperature *T_air_* [°C], (e) shear velocity *u*_∗_ [m s^−1^], (f) wind speed *u* [m s^−1^]. Colours represent different heights in the canopy (i.e. 3, 5, 7 & 8.5 m agl), solid lines depict times with more than 3 available observations and shade is standard deviation, dashed lines have 1-2 observations (due to repeated camera failure).

Leaf-to-air temperature differences (Δ*T_leaf_*_−*air*_, Fig. 3a, c) did not simply follow variations in SWR. For example, Δ*T_leaf_*_−*air*_ remained below 2.5 °C, and on average below 2 °C at 3 m and below 1.5 °C at 8.5 m in spite of similar levels of SWR (<550 W m^−2^). Variations in *T_leaf_* mostly paralleled air temperature (*T_air_*), but mostly remained above it (Fig. 3d). However, after 9:00 at 5 m and 7 m, the increase in *T_leaf_* slowed down below the rate of increase of *T_air_* (Fig. 3b, d), leading to a stabilisation or even decrease of Δ*T_leaf_*_−*air*_ even though SWR, *T_leaf_* and *T_air_* were still rising. In the exposed leaves at 7 m, this decrease even led to negative Δ*T_leaf_*_−*air*_ of up to −1 °C at the time of maximum *u*_∗_ and dropping SWR input around 15:00. Note that LWR at these exposed leaves was on average −40 W m^−2^ all day (i.e. radiative cooling). Against expectations, Δ*T_leaf_*_−*air*_ was highest at the bottom of the canopy (3 m) where it increased by ~1.5 °C following sunrise, in spite of low SWR (<100 W m^−2^). Furthermore, the independence of variations in *T_leaf_* from those in SWR was especially clear in this canopy layer (3 m), where the stabilisation of Δ*T_leaf_*_−*air*_ at ~2 °C (9:00-13:00) occurred before the peak of solar radiation (13:00-14:00).

Mean half-hourly wind speed (*u*) was not significantly different between different heights during daytime (At 11:15 and 15:45, *P* > .05; Fig. 3f). However, daytime turbulence (indicated using shear velocity *u*_∗_) did differ significantly between heights (*P* < .05), resulting from the interactions of wind speed and the canopy structure. For instance, *u*_∗_ was highest at the top of the canopy (0.5 m s^−1^; Fig. 3e). At 5, 7 & 8.5 m, the daily course of Δ*T_leaf_*_−*air*_ closely followed that of *T_air_* and turbulence: With *u*_∗_ < 0.25 m s^−1^, Δ*T_leaf_*_−*air*_ increased with *T_air_*, but quickly decreased above this threshold. At 3 m, *u*_∗_ remained below 0.25 m s^−1^ throughout the day, and Δ*T_leaf_*_−*air*_ remained high and appeared decoupled from *u*_∗_. In summary, our results do not show a direct dependence of Δ*T_leaf_*_−*air*_ on SWR and wind speed, but indicate more likely effects of that of the daily course of *T_air_* and *u*_∗_.

## Discussion

### Non-evaporative vs. evaporative cooling

Leaf thermal regulation has been shown across a wide range of ecosystems (Michaletz et al., 2016). It is commonly assumed that evaporative cooling through transpiration (TR) is the most effective heat dissipation mechanism (Wulfmeyer et al., 2014) and is key to optimize leaf temperature for the leaf biochemical processes (Dusenge et al., 2019). Evaporative cooling can result in a temperature reduction of up to 9 °C in broadleaf species (Urban et al., 2017). Furthermore, leaf temperature and Δ*T_leaf_*_−*air*_ provide an indicator of evaporative cooling, where warmer leaves indicate stomatal closure, more stress, and reduced gas exchange (e.g., Leuzinger and Körner, 2007; Kim et al., 2018). Evaporative cooling can also reflect the effects of other interacting factors that influence stomatal conductance, such as VPD (Kimball and Bernacchi, 2006), or rising CO_2_ levels (Urban et al., 2017; Kimball and Bernacchi, 2006; Leakey et al., 2006; Long et al., 2006). Similarly, the effect of climate driven rise in temperatures on leaf assimilation and water loss will depend on the effects of concurrent changes in factors such as VPD and atmospheric CO_2_ on leaf conductance, evaporation and, ultimately, leaf temperature (Dusenge et al., 2019; Kirschbaum and McMillan, 2018).

Our results indicated deviations from the common Δ*T_leaf_*_−*air*_ dependency on transpiration and stomatal conductance, as reflected in the similar Δ*T_leaf_*_−*air*_ in leaves with an order of magnitude difference in leaf transpiration. As discussed below, this can be achieved by changes in the leaf sensible heat flux, that must reflect changes in the leaf resistance to heat transport, which could be associated, in turn, with the small anatomical, structural and spectral differences observed between the leaves of drought-exposed and irrigated trees.

We observed that transpiration at the highest radiation bin (SWR ~800–900 W m^−2^, Fig. 2) translates to a latent heat LE of ~5 W m^−2^ in the drought-exposed, and 10× times higher value of ~103 W m^−2^ in the irrigated trees. This reflects a major change in the leaf energy budget, which can be described as:

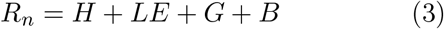

where *R_n_* is the net radiation, and *G* and *B* are net physical heat storage and energy taken up by biochemical reactions, which are typically considered negligible (Schymanski et al., 2013). *R_n_* in the two study plots was calculated from the net shortwave (using absorptance of 0.46; based on Hosgood et al., 1995) and longwave components 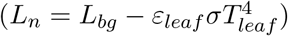. *R_n_* in the two study plots had similar mean values of 398 ± 8 W m^−2^ and 387 ± 17 W m^−2^ in the drought-exposed and irrigated plots, respectively, for the highest SWR bin (~800–900 W m^−2^, Fig. 2). With measured values of *R_n_* and LE, and negligible effects of *G* and *B* in Eq. 3, the leaf energy budget can only be adjusted via the sensible heat flux H. This, in turn, indicates a requirement for a large difference in H in the order of ~108 W m^−2^ between the drought-exposed (*H* ≈ 392W m^−2^) and irrigated (*H* ≈ 284W m^−2^) trees. The sensible heat flux can, in turn, be described by:

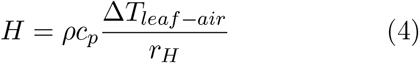

where *c_p_* and *ρ* are the heat capacity and density of air surrounding the leaves, and *r_H_* is the leaf resistance to heat transfer (where ‘leaf’ denotes multiple needles on a small twig; see Methods). Note that a simple sensitivity test indicates that a difference of 15 mmol mol^−1^ in air moisture or 5 °C in air temperature (when ambient *T_air_* is ~30°C) near leaves would produce a change in *H* of ~2.5– 10 W m^−2^ due to changes *c_p_* and *ρ*. Assuming, for a first approximation, that differences in air *c_p_* and *ρ* between the nearby plots were negligible, the difference in *H* derived above for the control and irrigated leaves, requires a proportional change in *r_H_*, (i.e. 392/284 = 1.38). In other words, under similar *R_n_* conditions, the efficient non-evaporative cooling observed in the control leaves must involve >30 % decrease in *r_H_*.

### Factors contributing to change in resistance to heat exchange at the leaf scale

Adaptation in plants has optimised leaf shapes and distribution for thermal regulation (Michaletz et al., 2016) and convective cooling (Smith, 1978). This can happen through an enhanced air flow and reduced *r_H_*, while concurrently minimizing its negative effects on evaporative water loss (Schymanski and Or, 2016; Grace and Wilson, 1976). Leaf and branch resistance to heat dissipation and the production of H is affected by factors that mainly modify the leaf boundary layer size or the flow of air surrounding the leaves. Often, *r_H_* has been parametrised using air flow variables measured above the canopy, i.e. wind speed u (Jones, 2004; Kool et al., 2016; Kustas and Norman, 1999; Verma, 1989) and shear velocity *u*_∗_ (Baldocchi and Ma, 2013; Liu et al., 2007), which indicates an increased local turbulence. Additionally, there are numerous reports of leaf anatomical, structural, and surface effects on convective cooling (Michaletz et al., 2016). For example, Leigh et al. (2012, 2017) demonstrate clear relationships between Δ*T_leaf_*_−*air*_ and leaf shape and thickness, resulting in a Δ*T_leaf_*_−*air*_ range of 2–10 °C. These studies also indicate the trade-offs between the leaf heat capacity that helps to deal with fast intense radiation exposure fluctuations (Schymanski et al., 2013), and the leaf size required to reduce *r_H_*. Leaf surface structures such as veins or hairs also affect roughness and therefore the generation of turbulence, which can improve convective cooling, as does leaf orientation relative to the wind direction (Schuepp, 1993; Grace and Wilson, 1976; Grace et al., 1980) or leaf spreading, as shown through the higher Δ*T_leaf_*_−*air*_ of densely packed needle-leaves (Smith and Carter, 1988). Needle-leaves, such as used in this study, may be a good example of low resistance leaves, based on their shape and surface to volume ratio, which all minimise the size of the boundary layer while maintaining sufficient leaf heat capacity.

Within the same species and at the same site, large changes in leaf and branch *r_H_* are more difficult to explain. But as argued above, the drought-exposed and irrigated trees must also exhibit some adjustments that produce large differences in *r_H_* between them. Indeed, preliminary observations indicated some changes in leaf/twig characteristics, resulting from the summer supplement irrigation. As a result, in the drought-exposed plot, needle-leaves were significantly shorter (and slightly but not significantly thicker), had a reduced clumpiness (needle density per twig), and lower chlorophyll content (Section S7, Figs. S5.1, S5.2 and S6.1). Shorter leaves and reduced packing are consistent with reduced Δ*T_leaf_*_−*air*_ (Smith and Carter, 1988) and thus *r_H_* in different plant species, as a result of simple vs. complex leaves that allow for more efficient air flow (Leigh et al., 2017). Early studies already noted the importance of leaf dimensions (e.g., Monteith, 1973; Landsberg and Thom, 1971; Gates, 1962 or packing of leaves (Michaletz and Johnson, 2006; Landsberg and Thom, 1971; Grant, 1984) in estimating the leaf boundary layer resistance to heat transfer *r_H_*. Hence, the observed change in leaf packing in our study (Section S7; Fig. S6.1) could indeed lead to a sufficient reduction in *r_H_* associated with changes in leaf boundary layer turbulence (Michaletz and Johnson, 2006). Additionally, the reduced pigment concentration under dry conditions (Section S7) could help in reducing leaf heating and increased reflectivity. Such effects could support the observed lack in Δ*T_leaf_*_−*air*_ differences, but this should be explored further in future work.

Clearly, the detailed estimation of changes in leaf resistance to heat dissipation requires, and justifies, further research. However, the available information provides strong supporting evidence to the hypothesis that significant changes in *r_H_* within the same species and location are possible under different soil moisture conditions. These conclusions, in turn, support the idea that efficient non-evaporative cooling can be developed in pine trees under dry conditions, based on a reduced *r_H_* and the generation of a large H. This results in a low Δ*T_leaf_*_−*air*_ similar to that of high transpiration leaves that rely on a combination of higher *r_H_* and higher TR to maintain a low Δ*T_leaf_*_−*air*_.

### Linking leaf temperature to canopy structure and turbulence

While the importance of surface roughness and structure is seldom considered at the leaf scale, such effects have been clearly demonstrated at the ecosystem scale (Rotenberg and Yakir, 2010, 2011). Sparse canopies, such as in our open-canopy forest, allow for the penetration of wind and efficient heat exchange through a higher roughness and reduced canopy-scale *r_H_* (Banerjee et al., 2017; Eder et al., 2015). In our conditions, this could reduce leaf temperatures to within 2 °C of the air. The vegetation type (conifers vs. broadleaves) and the observation point (top vs. within canopy) can also contribute to the range of observed canopy-air temperature differences of −2 °C to 10 °C (Table S7.1). Note that our use of non-aspirated radiation shields for air temperature measurements could lead to an overestimation of Δ*T_leaf_*_−*air*_ during daytime due to the effect of solar radiation, but the specific effect of such shields is likely to be small and with a similar effect in the different treatments. Such effects remain a matter of discussion in the literature (e.g., Kurzeja, 2009).

Our measurements of needle-leaf temperature in different heights of the canopy with distinct shading regimes showed that higher mean SWR does not always translate to a high Δ*T_leaf_*_−*air*_. Instead, *T_leaf_* generally paralleled *T_air_* during most of the day (Fig. 3b,d). In some periods, such as during the 9:00 to 14:00 period, the increase of *T_leaf_* fell below that of *T_air_*, hence Δ*T_leaf_*_−*air*_ no longer increased. Furthermore, at the bottom of the canopy (3 m agl) where SWR remained below 550 W m^−2^, Δ*T_leaf_*_−*air*_ reached 2 °C throughout the day, while in the exposed upper part of the canopy (7 m), Δ*T_leaf air_* dropped to ~0 °C in spite of high SWR of ~800 W m^−2^. Clearly, the results reflect complex interactions between *T_leaf_*, *T_air_* and SWR, but it also indicated a more tight relationship between *T_leaf_* and *T_air_* than with SWR. Note also that *T_leaf_* and *T_air_* at the bottom of the canopy are strongly affected by the soil temperature. In our example of a low density forest, SWR heats up a significant fraction of exposed ground, resulting in the production of large thermal radiation and sensible heat flux that heat up the bottom of the canopy, in what was previously described as a ‘canopy greenhouse effect’ (Rotenberg and Yakir, 2011).

In addition to the unexpectedly ambiguous effects of SWR on leaf temperature, the effects of wind speed and in turn, turbulence, can be at least as important. At the study site, wind speed was not directly correlated with Δ*T_leaf_*_−*air*_ (Fig. 3), but the decrease of Δ*T_leaf_*_−*air*_ with height coincided with the magnitude of turbulence, described using shear velocity *u*_∗_ (Fig. 3a,e). Indeed, with *u*_∗_ < 0.25m s^−1^, Δ*T_leaf_*_−*air*_ either increased or remained nearly constant, but decreased above this threshold. This explains the suppression of the expected high Δ*T_leaf_*_−*air*_ to ~0 °C at the top of the canopy (8.5 m), while it reached ~2 °C at the bottom (3 m) where turbulence was low (*u*_∗_ < 0.25m s^−1^).

These results provide an opportunity to explain the Δ*T_leaf_*_−*air*_ distribution across the canopy (Fig. 4), particularly in forests with a low stand density such as ours where turbulence creates an efficient air flow within the canopy layer. As noted above, at the bottom of the canopy (3 m agl) the combination of the lowest turbulence and the heating from the ground results in the highest Δ*T_leaf_*_−*air*_ of 2 °C (Fig. 3). Indeed, this ‘canopy greenhouse effect’ led to a comparable net radiation both above and within the canopy at the same research site (Rotenberg and Yakir, 2011), and therefore to a similar amount of heat to be dissipated. In exposed leaves at 7 m (mean 1⁄2h SWR: ~800 W m^−2^), Δ*T_leaf_−_air_* rapidly dropped from ~2 °C to ~0 °C by noon due to the high turbulence generated from the interaction of the typical regional sea breeze with the roughness of the forest canopy and in spite of a high air temperature. After 15:00, Δ*T_leaf_*_−*air*_ even dropped below ~0 °C in this height. It seems plausible that this is a result of the more dominant radiative cooling when incoming SWR sharply drops (negative net longwave radiation on average −40 W m^−2^ at this position), and slow decrease in TR at this time (see Qubaja et al., 2020). Such height dependence variations in the Δ*T_leaf_*_−*air*_ dynamics clearly reflect the complexity of the ambient conditions across the canopy under field conditions. Finally, Δ*T_leaf_*_−*air*_ of shaded leaves near the top of the canopy (8.5 m) was suppressed all day long and remained around ~1 °C. These leaves seemed to benefit from the combined effects of shading, the low incoming LWR from the cold atmosphere (as compared to LWR from the soil in the lower canopy), and maximal turbulence, which increases air-cooling through an increased turbulent air flow.

**Fig. 4.**
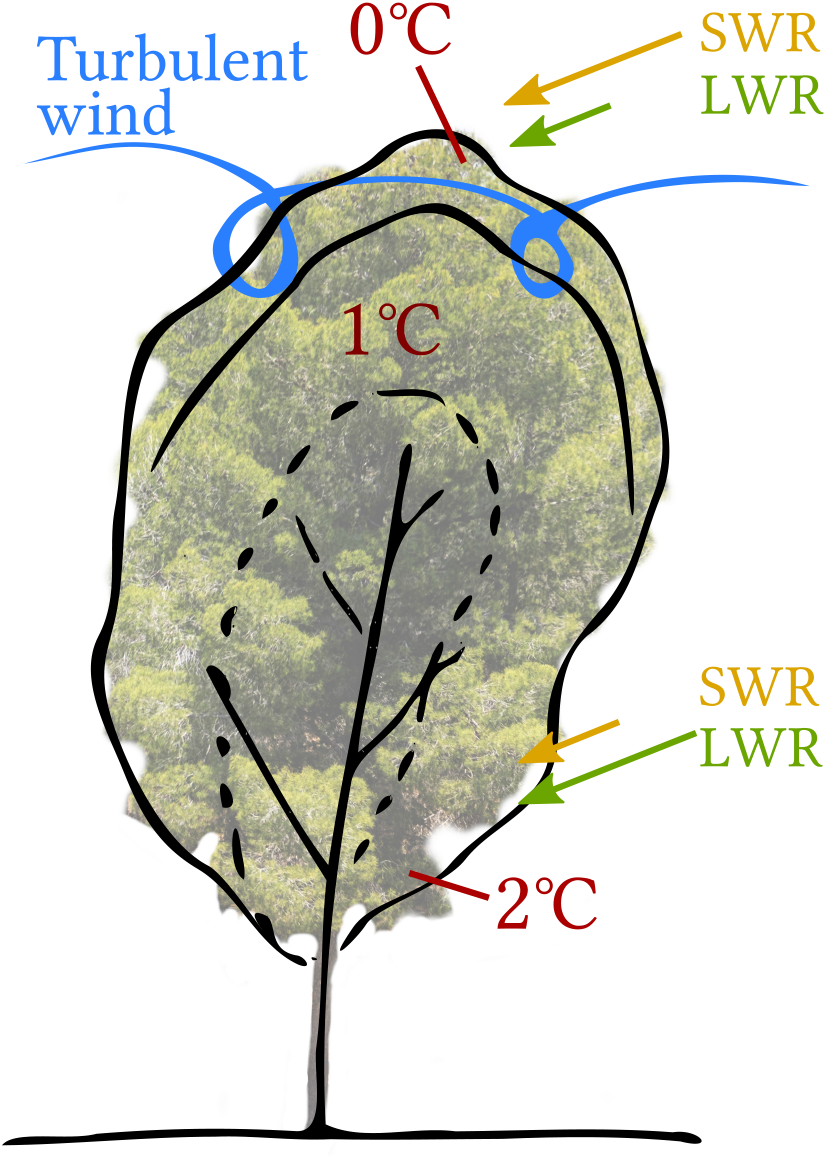
Conceptual diagram of the distribution of Δ*T_leaf_*_−*air*_ within the canopy structure, indicating the observed Δ*T_leaf_*_−*air*_ values in the different layers in this study, in relations to the radiation input of short wave and long wave radiations (SWR and LWR) from the surroundings as well as air cooling effects through turbulent wind.

The apparent contrast between the near-top and bottom of the canopy demonstrates the importance of turbulence within the canopy in controlling Δ*T_leaf_*_−*air*_, which relies on the reduced leaf-scale *r_H_* resulting from anatomical and biophysical adjustments. This efficient turbulent air cooling of leaves helps trees to survive the long summer drought (Tatarinov et al., 2016) and improves CO_2_ assimilation during the few optimal hours of the day (Maseyk et al., 2008; Michaletz et al., 2015). Indeed, leaf temperatures were kept below 40 °C at our site in high summer, which is considered the temperature of biochemical and physical damage to the photosynthetic apparatus (e.g., O’sullivan et al., 2017; Lancaster and Humphreys, 2020). The heat dissipated from leaves to their surrounding air can be subsequently transported to the atmosphere through turbulent wind penetration into the canopy layer. This has implications for the development of the large sensible heat flux out of the forest canopy, such as described by (Banerjee et al., 2017) as a basis of the generation of the so called canopy ‘convector effect’.

## Supporting information

Complete Supplement

## Acknowledgements

The authors are grateful to Tamir Dingjan for his help with needle-leaf identification in Python and for proof-reading, Huanhuan Wang for chlorophyll measurements, Irina Vishnevetsky for emissivity measurements of different materials and Revital Weic for her manual sampling of needle-leaf temperature.

This study was supported by by JNF-KKL (10-10-920-19) and a research grant from the *Yotam project* and the *Weizmann Institute Sustainability and Energy Research Initiative*. The long-term operation of the Yatir Forest Research Field Site is supported by the *Cathy Wills and Robert Lewis Program in Environmental Science*.

## Author contributions

The study was conceived by JM,ER and DY; JM developed the IR system,and carried out the measurements with the help of IO and FT; JM analysed the data under the guidance of ER and DY; JM,ER and DY contributed to the writing.

## Code and data availability

The data that support the findings of this study are available from the corresponding author upon reasonable request. Software scripts developed for the analysis are openly available as follows:

- Script to analyse gas exchange chamber flux data: ‘Branch-chamber-fluxes’ at https://doi.org/10.5281/zenodo.4284487, reference (Muller and Oz, 2020)
- Script to automatically trigger an infrared camera through a LAN interface: ‘FLIR-A320control’ at https://doi.org/10.5281/zenodo.4088156, reference (Muller, 2020)
- Python script to extract raw temperature data from FLIR infrared images: ‘IR-data-extraction’ at https://doi.org/10.5281/zenodo.4104314, reference (Muller and Segev, 2020)
- Script to detect pine needles and reference plates in infrared images: ‘Pine-needle-thermal-detection’ at https://doi.org/10.5281/zenodo.4284621, reference (Muller and Dingjan, 2020)

## Notes

https://doi.org/10.5281/zenodo.4088156

https://doi.org/10.5281/zenodo.4104314

https://doi.org/10.5281/zenodo.4284621

https://doi.org/10.5281/zenodo.4284487

### Competing Interest Statement

The authors have declared no competing interest.

### Summary of Updates

Sections and figures updated according to updated calculations

## References

Allen, C. D., Macalady, A. K., Chenchouni, H., Bachelet, D., McDowell, N., Vennetier, M., Kitzberger, T., Rigling, A., Breshears, D. D., Hogg, E. H. T., Gonzalez, P., Fensham, R., Zhang, Z., Castro, J., Demidova, N., Lim, J.-H., Allard, G., Running, S. W., Semerci, A., and Cobb, N. A global overview of drought and heat-induced tree mortality reveals emerging climate change risks for forests. Forest Ecology and Management, 259(4):660–684, February 2010. doi:10.1016/j.foreco.2009.09.001.

Allen, C. D., Breshears, D. D., and McDowell, N. On underestimation of global vulnerability to tree mortality and forest die-off from hotter drought in the Anthropocene. Ecosphere, 6(8):art129, 2015. doi:10.1890/ES15-00203.1.

Aubrecht, D. M., Helliker, B. R., Goulden, M. L., Roberts, D. A., Still, C., and Richardson, A. D. Continuous, long-term, high-frequency thermal imaging of vegetation: Uncertainties and recommended best practices. Agricultural and Forest Meteorology, 228–229:315–326, November 2016. doi:10.1016/j.agrformet.2016.07.017.

Baldocchi, D. and Ma, S. How will land use affect air temperature in the surface boundary layer? Lessons learned from a comparative study on the energy balance of an oak savanna and annual grassland in California, USA. Tellus B: Chemical and Physical Meteorology, 65(1):19994, December 2013. doi:10.3402/tellusb.v65i0.19994.

Baldocchi, D. and Penuelas, J. The physics and ecology of mining carbon dioxide from the atmosphere by ecosystems. Global Change Biology, 25 (4):1191–1197, 2019. doi:10.1111/gcb.14559.

Banerjee, T., De Roo, F., and Mauder, M. Explaining the convector effect in canopy turbulence by means of large-eddy simulation. Hydrology and Earth System Sciences (Online), 21(LA-UR-17-22651), 2017.

Birami, B., Gattmann, M., Heyer, A. G., Grote, R., Arneth, A., and Ruehr, N. K. Heat Waves Alter Carbon Allocation and Increase Mortality of Aleppo Pine Under Dry Conditions. Frontiers in Forests and Global Change, 1, 2018. doi:10.3389/ffgc.2018.00008.

Bonan, G. B. Forests and Climate Change: Forcings, Feedbacks, and the Climate Benefits of Forests. Science, 320(5882):1444–1449, June 2008. doi:10.1126/science.1155121.

Brugger, P., Banerjee, T., De Roo, F., Kröniger, K., Qubaja, R., Rohatyn, S., Rotenberg, E., Tatarinov, F., Yakir, D., Yang, F., and Mauder, M. Effect of Surface Heterogeneity on the Boundary-Layer Height: A Case Study at a Semi-Arid Forest. Boundary-Layer Meteorology, 169(2):233–250, November 2018. doi:10.1007/s10546-018-0371-5.

Drake, J. E., Tjoelker, M. G., V\a arhammar, A., Medlyn, B. E., Reich, P. B., Leigh, A., Pfautsch, S., Blackman, C. J., López, R., and Aspinwall, M. J. Trees tolerate an extreme heatwave via sustained transpirational cooling and increased leaf thermal tolerance. Global change biology, 2018.

Dusenge, M. E., Duarte, A. G., and Way, D. A. Plant carbon metabolism and climate change: elevated CO 2 and temperature impacts on photosynthesis, photorespiration and respiration. New Phytologist, 221(1):32–49, 2019.

Eder, F., De Roo, F., Rotenberg, E., Yakir, D., Schmid, H. P., and Mauder, M. Secondary circulations at a solitary forest surrounded by semi-arid shrubland and their impact on eddy-covariance measurements. Agricultural and Forest Meteorology, 211:115–127, 2015.

FLIR. FLIR A3xx series – User’s manual. Publ. No. T559498 Rev. a547, FLIR, July 2011.

Fritschen, L. J. and Gay, L. W. Environmental Instrumentation. Springer Science & Business Media, December 2012. ISBN 978-1-4612-6205-3.

Fuchs, M. Infrared measurement of canopy temperature and detection of plant water stress. Theoretical and Applied Climatology, 42(4):253–261, December 1990. doi:10.1007/BF00865986.

Gates, D. M. Leaf temperature and energy exchange. Archiv für Meteorologie, Geophysik und Bioklimatologie, Serie B, 12(2):321–336, October 1962. doi:10.1007/BF02315993.

Gates, D. M., Tibbals, E. C., and Kreith, F. Radiation and convection for ponderosa pine. American Journal of Botany, 52(1):66–71, 1965.

Geller, G. and Smith, W. Influence of leaf size, orientation, and arrangement on temperature and transpiration in three high-elevation, large-leafed herbs. Oecologia, 53(2):227–234, 1982. doi:10.1007/BF00545668.

Grace, J. and Wilson, J. The Boundary Layer over a Populus Leaf. Journal of Experimental Botany, 27 (2):231–241, April 1976. doi:10.1093/jxb/27.2.231.

Grace, J., Fasehun, F. E., and Dixon, M. Boundary layer conductance of the leaves of some tropical timber trees. Plant, Cell & Environment, 3(6): 443–450, 1980. doi:https://doi.org/10.1111/1365-3040.ep11586917.

Grant, R. H. The mutual interference of spruce canopy structural elements. Agricultural and Forest Meteorology, 32(2):145–156, August 1984. doi:10.1016/0168-1923(84)90084-4.

Gutschick, V. P. Leaf Energy Balance: Basics, and Modeling from Leaves to Canopies. In Hikosaka, K., Niinemets, [U+FFFD], and Anten, N. P., editors, Canopy Photosynthesis: From Basics to Applications, Advances in Photosynthesis and Respiration, pages 23–58. Springer Netherlands, Dordrecht, 2016. ISBN 978-94-017-7291-4. doi:10.1007/978-94-017-7291-4_2.

Hosgood, B., Jacquemoud, S., Andreoli, G., Verdebout, J., Pedrini, G., and Schmuck, G. Leaf optical properties experiment 93 (LOPEX93). Report EUR, 16095, 1995.

Idso, S. B., Jackson, R. D., Ehrler, W. L., and Mitchell, S. T. A Method for Determination of Infrared Emittance of Leaves. Ecology, 50(5):899–902, 1969. doi:10.2307/1933705.

Incropera, F. P., DeWitt, D. P., Bergman, T. L., and Lavine, A. S. Fundamentals of Heat and Mass Transfer. John Wiley & Sons, Hoboken, NJ, 6th edition edition, March 2006. ISBN 978-0-471-45728-2.

Jones, H. G. Application of Thermal Imaging and Infrared Sensing in Plant Physiology and Eco-physiology. In Advances in Botanical Research, volume 41 of Incorporating Advances in Plant Pathology, pages 107–163. Academic Press, January 2004. doi:10.1016/S0065-2296(04)41003-9.

Jones, H. G., Serraj, R., Loveys, B. R., Xiong, L., Wheaton, A., Price, A. H., Jones, H. G., Serraj, R., Loveys, B. R., Xiong, L., Wheaton, A., and Price, A. H. Thermal infrared imaging of crop canopies for the remote diagnosis and quantification of plant responses to water stress in the field. Functional Plant Biology, 36(11):978–989, November 2009. doi:10.1071/FP09123.

Kim, Y., Still, C., Hanson, C. V., Kwon, H., Greer, B. T., and Law, B. E. Canopy skin temperature variations in relation to climate, soil temperature, and carbon flux at a ponderosa pine forest in central Oregon. Agricultural and Forest Meteorology, 226–227:161–173, October 2016. doi:10.1016/j.agrformet.2016.06.001.

Kim, Y., Still, C., Roberts, D. A., and Goulden, M. L. Thermal infrared imaging of conifer leaf temperatures: Comparison to thermocouple measurements and assessment of environmental influences. Agricultural and Forest Meteorology, 248:361–371, January 2018. doi:10.1016/j.agrformet.2017.10.010.

Kimball, B. A. and Bernacchi, C. J. Evapotranspiration, Canopy Temperature, and Plant Water Relations. In Nösberger, J., Long, S. P., Norby, R. J., Stitt, M., Hendrey, G. R., and Blum, H., editors, Managed Ecosystems and CO2: Case Studies, Processes, and Perspectives, Ecological Studies, pages 311–324. Springer, Berlin, Heidelberg, 2006. ISBN 978-3-540-31237-6. doi:10.1007/3-540-31237-4_17.

Kirschbaum, M. U. F. and McMillan, A. M. S. Warming and Elevated CO2 Have Opposing Influences on Transpiration. Which is more Important? Current Forestry Reports, 4(2):51–71, June 2018. doi:10.1007/s40725-018-0073-8.

Kool, D., Kustas, W. P., Ben-Gal, A., Lazarovitch, N., Heitman, J. L., Sauer, T. J., and Agam, N. Energy and evapotranspiration partitioning in a desert vineyard. Agricultural and Forest Meteorology, 218:277–287, March 2016. doi:10.1016/j.agrformet.2016.01.002.

Kröniger, K., De Roo, F., Brugger, P., Huq, S., Banerjee, T., Zinsser, J., Rotenberg, E., Yakir, D., Rohatyn, S., and Mauder, M. Effect of Secondary Circulations on the Surface–Atmosphere Exchange of Energy at an Isolated Semi-arid Forest. Boundary-Layer Meteorology, 169(2):209–232, November 2018. doi:10.1007/s10546-018-0370-6.

Kurzeja, R. Accurate Temperature Measurements in a Naturally-Aspirated Radiation Shield. Boundary-Layer Meteorology, 134(1):181, November 2009. doi:10.1007/s10546-009-9430-2.

Kustas, W. P. and Norman, J. M. Evaluation of soil and vegetation heat flux predictions using a simple two-source model with radiometric temperatures for partial canopy cover. Agricultural and Forest Meteorology, 94(1):13–29, April 1999. doi:10.1016/S0168-1923(99)00005-2.

Lancaster, L. T. and Humphreys, A. M. Global variation in the thermal tolerances of plants. Proceedings of the National Academy of Sciences, 117(24):13580–13587, June 2020. doi:10.1073/pnas.1918162117.

Landsberg, J. J. and Thom, A. S. Aerodynamic properties of a plant of complex structure. Quarterly Journal of the Royal Meteorological Society, 97(414):565–570, 1971. doi:https://doi.org/10.1002/qj.49709741418.

Lapidot, O., Ignat, T., Rud, R., Rog, I., Alchanatis, V., and Klein, T. Use of thermal imaging to detect evaporative cooling in coniferous and broadleaved tree species of the Mediterranean maquis. Agricultural and Forest Meteorology, 271:285–294, June 2019. doi:10.1016/j.agrformet.2019.02.014.

Leakey, A. D., Uribelarrea, M., Ainsworth, E. A., Naidu, S. L., Rogers, A., Ort, D. R., and Long, S. P. Photosynthesis, productivity, and yield of maize are not affected by open-air elevation of CO2 concentration in the absence of drought. Plant physiology, 140(2):779–790, 2006.

Leigh, A., Sevanto, S., Ball, M. C., Close, J. D., Ellsworth, D. S., Knight, C. A., Nicotra, A. B., and Vogel, S. Do thick leaves avoid thermal damage in critically low wind speeds? New Phytologist, 194(2): 477–487, 2012. doi:https://doi.org/10.1111/j.1469-8137.2012.04058.x.

Leigh, A., Sevanto, S., Close, J. D., and Nicotra, A. B. The influence of leaf size and shape on leaf thermal dynamics: does theory hold up under natural conditions? Plant, cell & environment, 40 (2):237–248, 2017.

Leuzinger, S. and Körner, C. Tree species diversity affects canopy leaf temperatures in a mature temperate forest. Agricultural and Forest Meteorology, 146(1):29–37, September 2007. doi:10.1016/j.agrformet.2007.05.007.

Leuzinger, S., Vogt, R., and Körner, C. Tree surface temperature in an urban environment. Agricultural and Forest Meteorology, 150(1):56–62, January 2010. doi:10.1016/j.agrformet.2009.08.006.

Liu, S., Lu, L., Mao, D., and Jia, L. Evaluating parameterizations of aerodynamic resistance to heat transfer using field measurements. Hydrology and Earth System Sciences Discussions, 11(2):769–783, 2007.

Long, S. P., Humphries, S., and Falkowski, P. G. Photoinhibition of photosynthesis in nature. Annual review of plant biology, 45(1):633–662, 1994.

Long, S. P., Ainsworth, E. A., Leakey, A. D. B., Nösberger, J., and Ort, D. R. Food for Thought: Lower-Than-Expected Crop Yield Stimulation with Rising CO2 Concentrations. Science, 312(5782):1918–1921, June 2006. doi:10.1126/science.1114722.

Maimaitijiang, M., Sagan, V., Sidike, P., Hartling, S., Esposito, F., and Fritschi, F. B. Soybean yield prediction from UAV using multimodal data fusion and deep learning. Remote Sensing of Environment, 237:111599, February 2020. doi:10.1016/j.rse.2019.111599.

Maseyk, K. S. Ecophysiological and phenological aspects of Pinus halepensis in an arid-Mediterranean environment. Ph.D., The Weizmann Institute of Science (Israel), Israel, 2006. URL https://search.proquest.com/docview/304955792/citation/84F5442BDD884B15PQ/1.

Maseyk, K. S., Lin, T., Rotenberg, E., Grünzweig, J. M., Schwartz, A., and Yakir, D. Physiology–phenology interactions in a productive semi-arid pine forest. New Phytologist, 178(3): 603–616, 2008. doi:https://doi.org/10.1111/j.1469-8137.2008.02391.x.

McMahon, T. A., Peel, M. C., Lowe, L., Srikanthan, R., and McVicar, T. R. Estimating actual, potential, reference crop and pan evaporation using standard meteorological data: a pragmatic synthesis. Hydrol. Earth Syst. Sci, 17(4):1331–1363, 2013.

Michaletz, S. T. and Johnson, E. A. Foliage influences forced convection heat transfer in conifer branches and buds. New Phytologist, 170(1):87–98, 2006. doi:10.1111/j.1469-8137.2006.01661.x.

Michaletz, S. T., Weiser, M. D., Zhou, J., Kaspari, M., Helliker, B. R., and Enquist, B. J. Plant Thermoregulation: Energetics, Trait–Environment Interactions, and Carbon Economics. Trends in Ecology & Evolution, 30(12):714–724, December 2015. doi:10.1016/j.tree.2015.09.006.

Michaletz, S. T., Weiser, M. D., McDowell, N. G., Zhou, J., Kaspari, M., Helliker, B. R., and Enquist, B. J. The energetic and carbon economic origins of leaf thermoregulation. Nature Plants, 2(9):1–9, August 2016. doi:10.1038/nplants.2016.129.

Monteith, J. L. Principles of environmental physics Edward Arnold. London, 214p, 1973.

Muller, J. D. FLIR-A320-control: Tool to remotely focus and trigger a FLIR A320 infrared camera, October 2020. URL https://zenodo.org/record/4088156.

Muller, J. D. and Dingjan, T. Pine-needle-thermal-detection: Tool to detect pine needles in thermal images, November 2020. URL https://zenodo.org/record/4284621.

Muller, J. D. and Oz, I. Branch-chamber-fluxes: Branch-chamber-fluxes, November 2020. URL https://zenodo.org/record/4284487.

Muller, J. D. and Segev, L. IR-data-extraction: Tool to extract raw temperature data from FLIR R-jpegs, October 2020. URL https://zenodo.org/record/4104314.

Muller, J. D., Rotenberg, E., Tatarinov, F., Vishnevetsky, I., Dingjan, T., Kribus, A., and Yakir, D. Dual reference method for high precision infrared measurement of leaf surface temperature under field conditions. bioRxiv, page 2021.04.25.440729, April 2021. doi:10.1101/2021.04.25.440729.

Neriah, A. B., Assouline, S., Shavit, U., and Weisbrod, N. Impact of ambient conditions on evaporation from porous media. Water Resources Research, 50(8):6696–6712, 2014. doi:https://doi.org/10.1002/2014WR015523.

O’sullivan, O. S., Heskel, M. A., Reich, P. B., Tjoelker, M. G., Weerasinghe, L. K., Penillard, A., Zhu, L., Egerton, J. J. G., Bloomfield, K. J., Creek, D., Bahar, N. H. A., Griffin, K. L., Hurry, V., Meir, P., Turnbull, M. H., and Atkin, O. K. Thermal limits of leaf metabolism across biomes. Global Change Biology, 23(1):209–223, January 2017. doi:10.1111/gcb.13477.

Pau, S., Detto, M., Kim, Y., and Still, C. Tropical forest temperature thresholds for gross primary productivity. Ecosphere, 9(7):e02311, 2018. doi:https://doi.org/10.1002/ecs2.2311.

Peel, M. C., Finlayson, B. L., and McMahon, T. A. Updated world map of the Köppen-Geiger climate classification. Hydrology and Earth System Sciences, 11(5):1633–1644, October 2007. doi:10.5194/hess-11-1633-2007.

Preisler, Y. Water-use strategies leading to resilience of pine trees to global climatic change. PhD Thesis, Hebrew University of Jerusalem, Rehovot, Israel, December 2019.

Qubaja, R., Grünzweig, J. M., Rotenberg, E., and Yakir, D. Evidence for large carbon sink and long residence time in semiarid forests based on 15 year flux and inventory records. Global Change Biology, 2019. doi:10.1111/gcb.14927.

Qubaja, R., Amer, M., Tatarinov, F., Rotenberg, E., Preisler, Y., Sprintsin, M., and Yakir, D. Partitioning evapotranspiration and its long-term evolution in a dry pine forest using measurement-based estimates of soil evaporation. Agricultural and Forest Meteorology, 281:107831, February 2020. doi:10.1016/j.agrformet.2019.107831.

Richardson, A. D., Aubrecht, D. M., Basler, D., Hufkens, K., Muir, C. D., and Hanssen, L. Developmental changes in the reflectance spectra of temperate deciduous tree leaves, and implications for thermal emissivity and leaf temperature. New Phytologist, n/a(n/a), 2020. doi:10.1111/nph.16909.

Rotenberg, E. and Yakir, D. Contribution of Semi-Arid Forests to the Climate System. Science, 327(5964):451–454, January 2010. doi:10.1126/science.1179998.

Rotenberg, E. and Yakir, D. Distinct patterns of changes in surface energy budget associated with forestation in the semiarid region. Global Change Biology, 17(4):1536–1548, 2011. doi:10.1111/j.1365-2486.2010.02320.x.

Schuepp, P. H. Tansley Review No. 59. Leaf Boundary Layers. The New Phytologist, 125(3):477–507, 1993.

Schymanski, S. J. and Or, D. Wind increases leaf water use efficiency. Plant, Cell & Environment, 39(7):1448–1459, 2016. doi:10.1111/pce.12700.

Schymanski, S. J., Or, D., and Zwieniecki, M. Stomatal Control and Leaf Thermal and Hydraulic Capacitances under Rapid Environmental Fluctuations. PLOS ONE, 8(1):e54231, January 2013. doi:10.1371/journal.pone.0054231.

Siebert, S., Ewert, F., Rezaei, E. E., Kage, H., and Graß, R. Impact of heat stress on crop yield—on the importance of considering canopy temperature. Environmental Research Letters, 9(4):044012, April 2014. doi:10.1088/1748-9326/9/4/044012.

Smith, W. K. Temperatures of Desert Plants: Another Perspective on the Adaptability of Leaf Size. Science, 201(4356):614–616, August 1978. doi:10.1126/science.201.4356.614.

Smith, W. K. and Carter, G. A. Shoot structural effects on needle temperatures and photosynthesis in conifers. American Journal of Botany, 75(4): 496–500, 1988.

Smith, W. K., Dannenberg, M. P., Yan, D., Herrmann, S., Barnes, M. L., Barron-Gafford, G. A., Biederman, J. A., Ferrenberg, S., Fox, A. M., Hudson, A., Knowles, J. F., MacBean, N., Moore, D. J. P., Nagler, P. L., Reed, S. C., Rutherford, W. A., Scott, R. L., Wang, X., and Yang, J. Remote sensing of dryland ecosystem structure and function: Progress, challenges, and opportunities. Remote Sensing of Environment, 233:111401, November 2019. doi:10.1016/j.rse.2019.111401.

Song, Q.-H., Deng, Y., Zhang, Y. P., Deng, X.-B., Lin, Y.-X., Zhou, L.-G., Fei, X.-H., Sha, L.-Q., Liu, Y.-T., Zhou, W.-J., and Gao, J.-B. Comparison of infrared canopy temperature in a rubber plantation and tropical rain forest. International Journal of Biometeorology, 61(10):1885–1892, October 2017. doi:10.1007/s00484-017-1375-4.

Still, C., Powell, R., Aubrecht, D., Kim, Y., Helliker, B., Roberts, D., Richardson, A. D., and Goulden, M. Thermal imaging in plant and ecosystem ecology: applications and challenges. Ecosphere, 10 (6), 2019.

Still, C., Rastogi, B., Page, G. F. M., Griffith, D. M., Sibley, A., Schulze, M., Hawkins, L., Pau, S., Detto, M., and Helliker, B. R. Imaging canopy temperature: shedding (thermal) light on ecosystem processes. New Phytologist, n/a(n/a), 2021. doi:10.1111/nph.17321.

Tatarinov, F., Rotenberg, E., Maseyk, K., Ogée, J., Klein, T., and Yakir, D. Resilience to seasonal heat wave episodes in a Mediterranean pine forest. New Phytologist, 210(2):485–496, April 2016. doi:10.1111/nph.13791.

Tattersall, G. Thermimage, v.4.1.0, December 2019. URL https://zenodo.org/record/3590036.

Taylor, S. E. Optimal Leaf Form. In Gates, D. M. and Schmerl, R. B., editors, Perspectives of Biophysical Ecology, Ecological Studies, pages 73–86. Springer, Berlin, Heidelberg, 1975. ISBN 978-3-642-87810-7. doi:10.1007/978-3-642-87810-7_5.

Tibbals, E. C., Carr, E. K., Gates, D. M., and Kreith, F. Radiation and convection in conifers. American Journal of Botany, 51(5):529–538, 1964.

Urban, J., Ingwers, M. W., McGuire, M. A., and Teskey, R. O. Increase in leaf temperature opens stomata and decouples net photosynthesis from stomatal conductance in Pinus taeda and Populus deltoides x nigra. Journal of Experimental Botany, 68(7):1757–1767, March 2017. doi:10.1093/jxb/erx052.

Verma, S. B. Aerodynamic resistances to transfers of heat, mass and momentum. Estimation of areal evapotranspiration, 177:13–20, 1989.

Virtanen, P., Gommers, R., Oliphant, T. E., Haberland, M., Reddy, T., Cournapeau, D., Burovski, E., Peterson, P., Weckesser, W., Bright, J., van der Walt, S. J., Brett, M., Wilson, J., Millman, K. J., Mayorov, N., Nelson, A. R. J., Jones, E., Kern, R., Larson, E., Carey, C. J., Polat, I., Feng, Y., Moore, E. W., VanderPlas, J., Laxalde, D., Perktold, J., Cimrman, R., Henriksen, I., Quintero, E. A., Harris, C. R., Archibald, A. M., Ribeiro, A. H., Pedregosa, F., and van Mulbregt, P. SciPy 1.0: fundamental algorithms for scientific computing in Python. Nature Methods, 17(3):261–272, March 2020. doi:10.1038/s41592-019-0686-2.

Vishnevetsky, I., Rotenberg, E., Kribus, A., and Yakir, D. Method for accurate measurement of infrared emissivity for opaque low-reflectance materials. Applied Optics, 58(17):4599–4609, 2019.

Werner, C., Correia, O., Beyschlag, W., Werner, C., Correia, O., and Beyschlag, W. Characteristic patterns of chronic and dynamic photoinhibition of different functional groups in a Mediterranean ecosystem, Characteristic patterns of chronic and dynamic photoinhibition of different functional groups in a Mediterranean ecosystem. Functional Plant Biology, Functional Plant Biology, 29, 29(8, 8):999, 999–1011, 1011, August 2002. doi:10.1071/PP01143, 10.1071/PP01143.

Wulfmeyer, V., Branch, O., Warrach-Sagi, K., Bauer, H.-S., Schwitalla, T., and Becker, K. The Impact of Plantations on Weather and Climate in Coastal Desert Regions. Journal of Applied Meteorology and Climatology, 53(5):1143–1169, January 2014. doi:10.1175/JAMC-D-13-0208.1.

Yi, K., Smith, J. W., Jablonski, A. D., Tatham, E. A., Scanlon, T. M., Lerdau, M. T., Novick, K. A., and Yang, X. High Heterogeneity in Canopy Temperature Among Co-occurring Tree Species in a Temperate Forest. Journal of Geophysical Research: Biogeosciences, 125(12):e2020JG005892, 2020. doi:https://doi.org/10.1029/2020JG005892.

Zhang, L., Niu, Y., Zhang, H., Han, W., Li, G., Tang, J., and Peng, X. Maize Canopy Temperature Extracted From UAV Thermal and RGB Imagery and Its Application in Water Stress Monitoring. Frontiers in Plant Science, 10, 2019. doi:10.3389/fpls.2019.01270.

