## Supplementary material for "Evidence for efficient non-evaporative leaf-to-air heat dissipation in a pine forest under drought conditions": Complete Supplement

### Supporting Information

#### S1 Automatic sampling of leaf temperatures

**Technical summary of leaf identification:** A matrix of raw infrared temperatures were extracted from the FLIR infrared images using a Python script developed based on an R script by Tattersall (2019). Different areas of interest were scanned in horizontal lines to identify needle leaves and reference plates. A peak detection algorithm contained in SciPy (Virtanen et al., 2020) was used on each line to create a mask of valid pixels of each category: (a) To identify needles, a peak prominence of a height of ⅓ of the total range of values in each line and a maximum width of 5 px was used; (b) to identify reference plates, the junction of values between the 'warm' emissive and the 'cold' reflective plate was identified by calculating the first discrete difference along the horizontal line of data points, areas with a fixed width of 20 px around that junction were selected and peaks in those areas were removed (e.g. for leaves in front of the plates). The median, mean and standard deviations of all leaf temperatures of a twig were calculated for all the pixels in each category of data (i.e. leaves and reference plates), but medians were used to reduce the effect of obvious outliers.

**Comparison to manual sampling:** The results of this script were compared to manual sampling of leaf temperatures during the initial month of measurements ($R^2 = 0.99$, $P < .001$, $RMSE = 0.13$; Fig. S1.1), which confirms the robustness of our script. Then, $\frac{1}{2}$h means and standard deviations were calculated from the median and standard deviations of leaf temperatures of each infrared image. Manual sampling was not as accurate because the hand-drawn polygons were often more than 5 pixels wide, which introduces small errors, thus explaining the variation of measurements: At high temperatures (daytime), the manually sampled polygons captured a larger amount of the hot background (mostly soil). In lower temperatures (night), the difference between leaves and background was smaller, thus explaining the smaller divergence in values. Thus, manual sampling tended to overestimate leaf temperature.

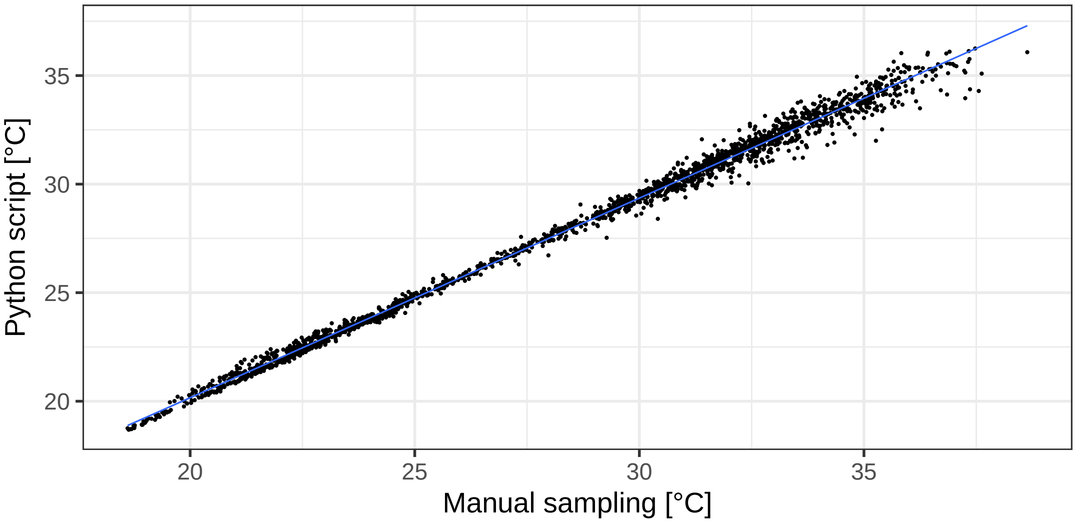


**Fig. S1.1.** Comparison between manual sampling of leaf temperature (by Revital Weic) and the automatic python script, where $y = 1.08x - 1.76$, $R^2 = 0.99$, $P < .001$. Manual sampling typically included polygons more than 5 pixels wide, which introduces small errors

#### S2 Effect of solar radiation on air temperature measurements

We built a simple levelled calibration stand on a tripod (Fig. S2.1), which was deployed outdoors for 2 days.

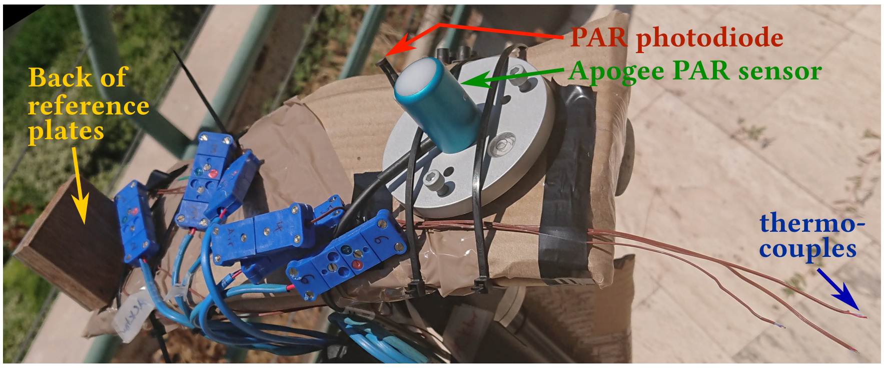


**Fig. S2.1.** Calibration setup, showing the Apogee SQ-500-ss PAR sensor (green), Hamamatsu G1118 PAR photodiode (red), IR reference plates (yellow), air temperature thermocouple (blue) and GA10K3MCD1 air temperature thermistor (not visible)

The Hamamatsu G1118 photodiode (Hamamatsu photonics K.K., Iwata City, Japan) was calibrated against the Apogee SQ-500-ss PAR sensor (Agogee Instruments Inc., Logan UT, USA; Fig. S2.2). Due to its lack of a diffusing filter, the photodiode was much more sensitive to levelling, resulting in a slight hysteresis shown as a result of solar angle. A high correlation coefficient ($y = 20.27 + 1.87x$, $R^2 = 0.99$, $P < .001$) indicates that this hysteresis has a minor effect, however.

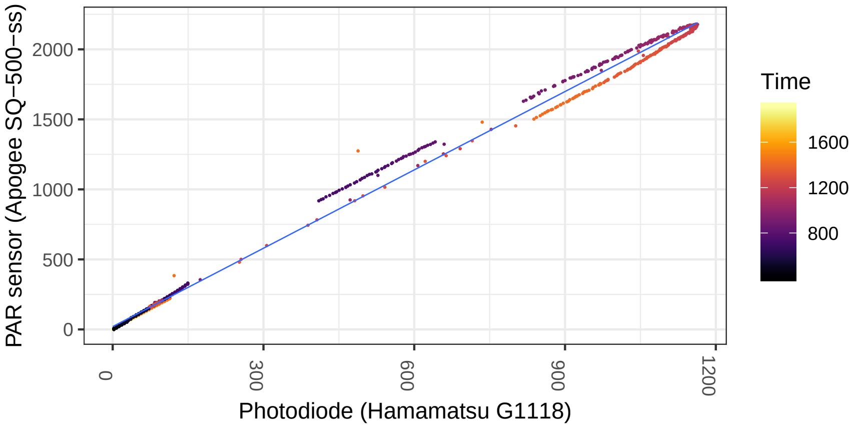


**Fig. S2.2.** Measurements of photosynthetically active radiation (PAR; µmol mol$^{-2}$ s$^{-1}$) by the photodiode (Hamamatsu G1118) and the professional-grade PAR sensor (Apogee SQ-500-ss). The hysteresis shows the sensitivity of the photodiode to levelling and results from a changing solar angle (shown as a colour using time). Linear correlation (blue): $y = 20.27 + 1.87x$, $R^2 = 0.99$, $P < .001$

Then, the air temperature thermocouple was correlated with the trusted thermistor (GA10K3MCD1; TE connectivity Ltd., Schaffhausen, Switzerland) during the night to avoid the effect of solar radiation on the measurements (Fig. S2.3). The measurements show a near 1:1 correlation ($y = 0.21 + 0.99x$, $R^2 = 0.99$, $P < .001$), meaning that a further calibration of the thermocouple was unnecessary.

Finally, we assessed the effect of fast-changing solar radiation on the temperature of the reference plates used in the infrared camera system. As expected, the plates heated

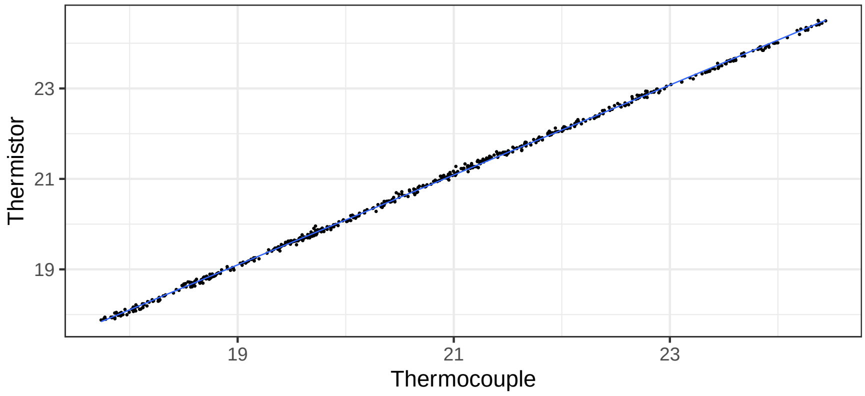


**Fig. S2.3.** Thermocouple vs Thermistor air temperature measurements at night, when the wind speed is minimal, thus removing noisy small-scale fluctuations. Correlation: $y = 0.21 + 0.99x$, $R^2 = 0.99$, $P < .001$

up or cooled down quickly due to a change in PAR (Fig. S2.4). These changes do not, however, affect the calibration of the infrared camera

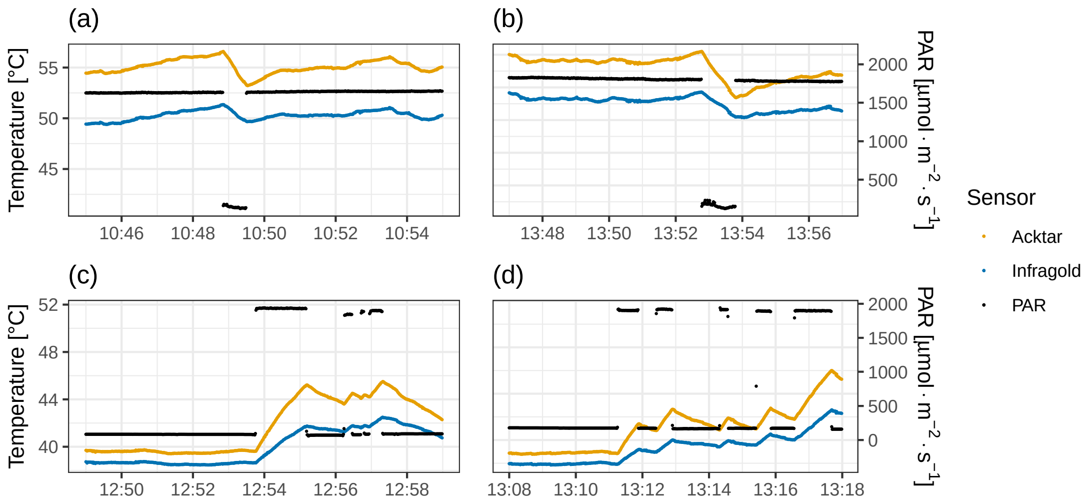


**Fig. S2.4.** Four events showing the effect of fast changes of photosynthetically active radiation (PAR, black) on the temperature of the emissive reference plate (Acktar coating, yellow) and the infragold coating (blue).

#### S3 Independent measurement of leaf-air temperature difference

Additional measurements of $\Delta T_{leaf-air}$ were made with a Mini-PAM II sensor's open system exposed to ambient light (Heinz Walz GmbH, Effeltrich, Germany) that employs a leaf thermocouple (Fig. S3.1). No significant difference between drought-exposed (control) and irrigated trees was detected.

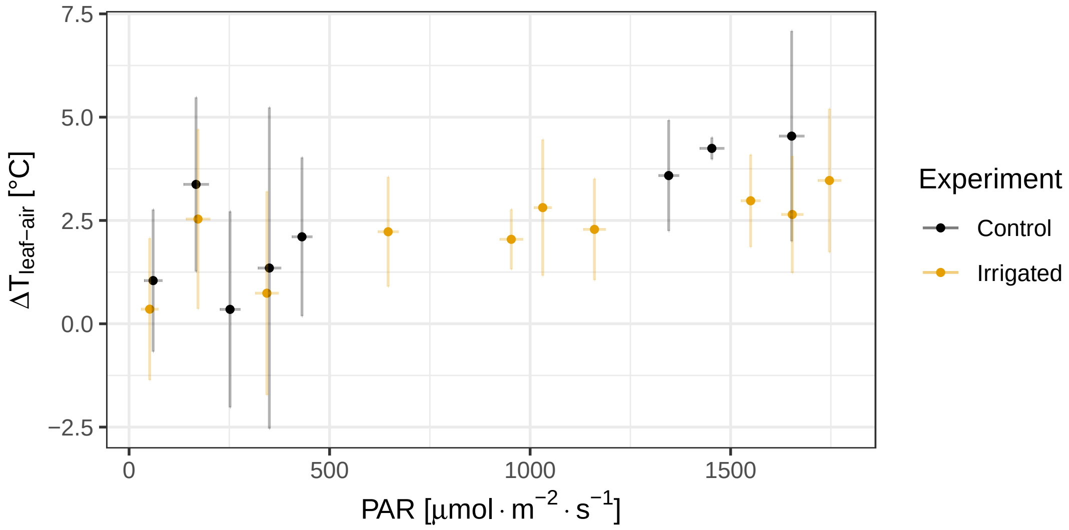


**Fig. S3.1.** Relationship between leaf-to-air temperature difference $\Delta T_{leaf-air}$ and photosynthetically active radiation PAR in bins of 100 µmol m$^{-2}$ s$^{-1}$, measured approximately monthly from June 2019 to July 2020 in irrigated (yellow) and drought-exposed trees (black). Bars represent standard deviations

#### S4 Measurement locations within the site

Figure S4.1 shows the Yatir forest research site with all the measurement locations used in the framework of this research. Data presented in this paper includes measurements in (2a) an experimentally irrigated and (2b) a drought-exposed plot, as well as (3) measurements in different heights throughout the canopy. Dates and times for measurements with the sonic anemometers (turbulence and wind measurements) are summarised in Table S4.1.

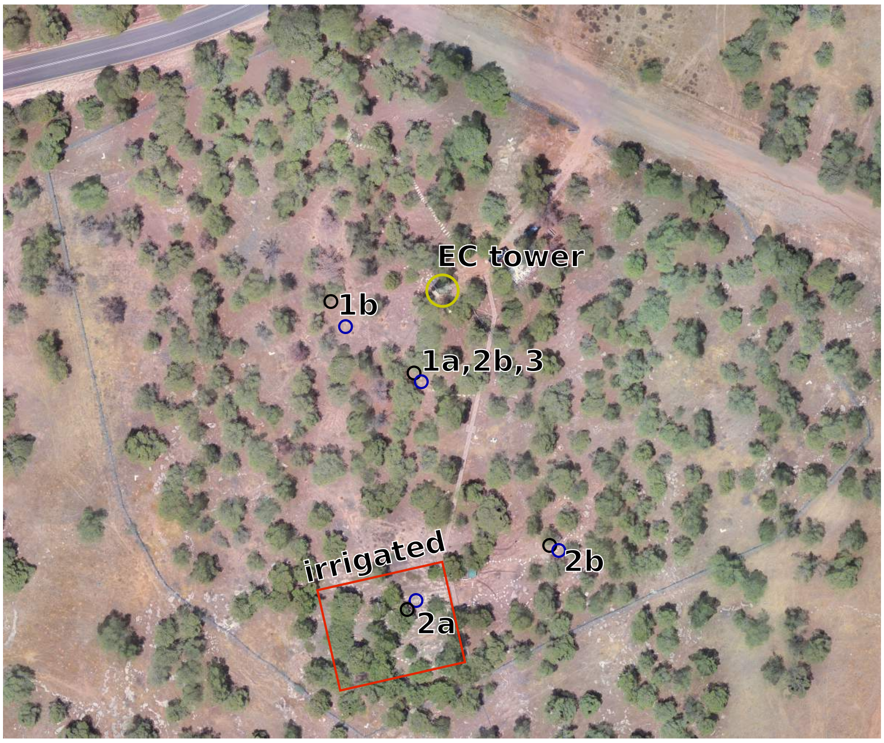


**Fig. S4.1.** Aerial map of the Yatir research site (2020-06-25; DJI Matrice 200 with DJI Zenmuse XT2 camera, processed with Pix4Dmapper). The yellow circle represents the Eddy Covariance (EC) flux tower. Black circles are the locations of the infrared camera mast setup; blue circles are locations of the sonic anemometer mast used for wind speed measurements. Experiments are numbered: (1) Long-term measurements across seasons (summer 2017-autumn 2019; infrared camera in 5 m from June 2018; sonic anemometers in 3 and 10 m), in (a) a dense canopy and (b) a gap in the canopy; (2) Setup (summers 2018 & 2019; infrared camera setup in 5 m) with simultaneous gas-exchange measurements in (a) an experimentally irrigated plot and (b) under drought-exposed (control) conditions; and (3) measurements in different heights (summer 2018; 3, 5, 7 & 8.5 m). Unless specified, the sonic anemometers were installed close together (1.5 m distance from each other) to measure the slice of the atmosphere in which the infrared camera setup was located.

**Table S4.1.** Setup of instrumentation masts and dates of measurements according to sets of measurements: (a) drought-exposed and irrigated plots and (b) measurements in slices

| Experiment | Height of IR camera | Height of sonics | Dates |
|---|---|---|---|
| Irrigation / drought-exposed | | | |
| Drought 2018 | 5m | 435cm & 565cm | 2018-07-01 to 2018-07-05; 2018-07-20 to 2018-07-30 |
| Irrigation 2018 | 4.75m | 410cm & 540cm | 2018-08-29 to 2018-10-30 |
| Drought 2019 | 5m | 435cm & 565cm | 2019-07-31 to 2019-09-14 |
| Irrigation 2019 | 5.75m | 510cm & 640cm | 2019-06-25 to 2019-07-31 |
| Canopy slices in different heights | | | |
| 3m | 3m | 240cm & 360cm | 2018-07-05 to 2018-07-19 |
| 5m | 5m | 435cm & 565cm | 2018-07-20 to 2018-07-30 |
| 7m | 7m | 635cm & 765cm | 2018-08-14 to 2018-08-28 |
| 8.5m | 8.5m | 775cm & 905cm | 2018-08-01 to 2018-08-13 |

#### S5 Leaf dimension differences between drought-exposed and irrigated trees

Needle-leaf dimensions were regularly measured monthly at our site in both drought-exposed and irrigated plots, before and after the onset of irrigation in May 2017. Irrigation led to a rapid leaf elongation, but length differences remained significant even after the first year (Fig. S5.1). Differences in needle-leaf diameters were smaller than the error of the measurement (Fig. S5.2). A summary of the differences is given in Section S7.

**Table S5.1.** Factors that could affect $r_H$: needle-leaf dimensions (Figs. S5.1 and S5.2), summer chlorophyll content and clumpiness of leaves, i.e. the ratio between the projected (Fig. S6.1) and total leaf area

| Treatment | Drought-exposed | Irrigated |
|---|---|---|
| Leaf length [mm] | 4.58±0.70 | 8.27±0.40 |
| Leaf diameter [mm] | 0.84±0.04 | 0.82±0.02 |
| Chlorophyll content [$\mu g\,cm^{-2}$] | 34.31±4.61 | 47.97±8.58 |
| Clumpiness [$m^2\,m^{-2}$] | 3.18±0.016 | 6.16±0.077 |

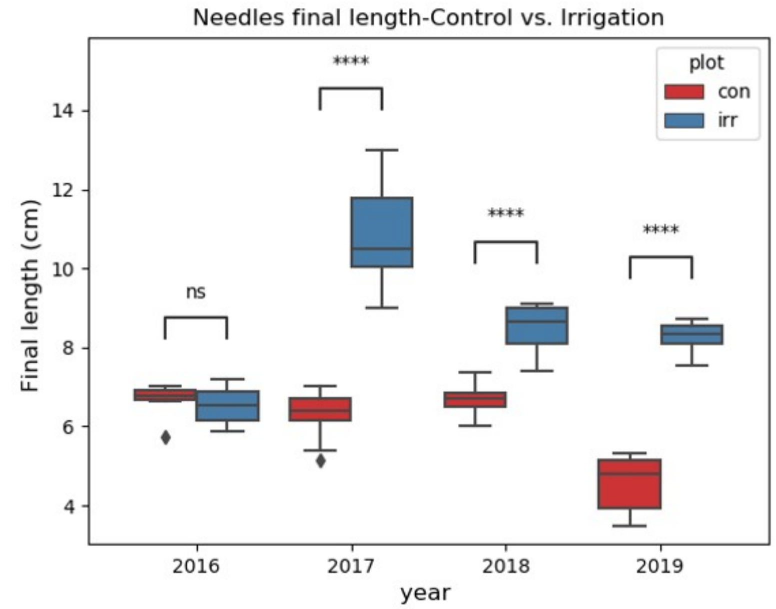


**Fig. S5.1.** Needle-leaf length comparisons between irrigated (irr) and drought-exposed control (con) plots for each year of measurements. Irrigation began in May 2017

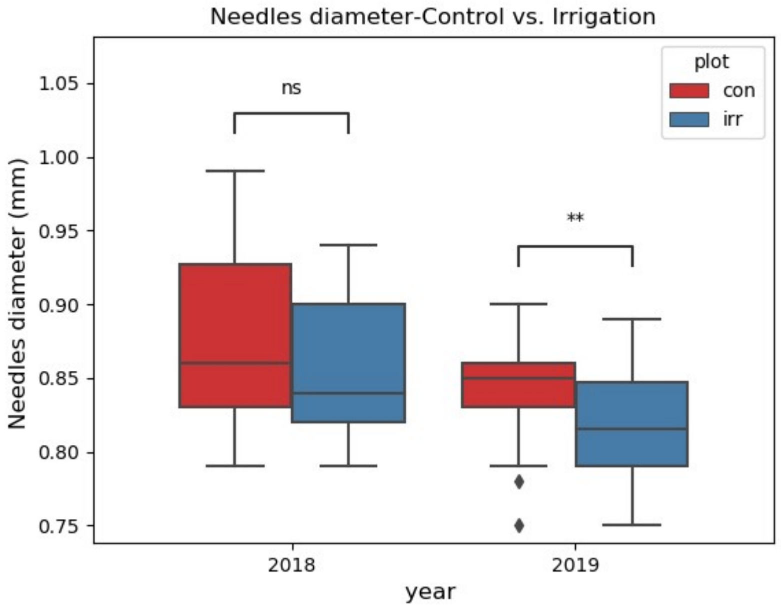


**Fig. S5.2.** Needle-leaf diameter comparisons between irrigated (irr) and drought-exposed control (con) plots for 2018-2019, with a measurement error of >0.1mm

#### S6 Needle-leaf distribution in drought-exposed and irrigated trees

The spread of needle-leaves was qualitatively assessed on a branch of an irrigated and drought-exposed tree, respectively. Leaves seemed to be more clumped on irrigated trees, which is assumed to reduce air flow (Fig. S6.1).

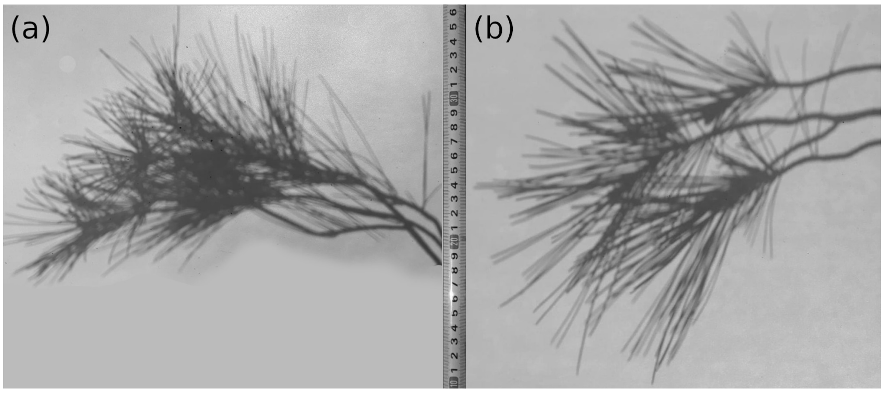


**Fig. S6.1.** Needle-leaf distribution on a branch of an (a) irrigated and (b) drought-exposed tree

#### S7 Canopy-to-air temperature differences (literature review)

Data of leaf- and/or canopy-to-air temperature differences $\Delta T_{canopy-air}$ measured with infrared thermographers was collected from multiple non-exhaustive literature sources, either directly available in the text or extracted from graphs. In some cases, when leaf or canopy and air temperatures were available separately, $\Delta T_{canopy-air}$ was calculated here rather than providing the two values separately. In most studies, air temperature was measured at nearby meteorological stations, except for Drake et al. (2018) who measured it inside their experimental setup. The data, summarised in Table S7.1, shows that $\Delta T_{canopy-air}$ is much higher in closed canopy forests than in free-standing trees.

The Köppen-Geiger categories mentioned in Table S7.1 are the following:

| Köppen-Geiger category | Descriptive category |
|---|---|
| *Am* | Tropical monsoon climate |
| *Cfa* | Humid subtropical climate |
| *Cfb* | Oceanic climate |
| *Csa* | Hot-summer Mediterranean climate |
| *Csb* | Warm-summer Mediterranean climate |
| *Dfb* | Warm summer continental climate |

**Table S7.1.** Leaf/Canopy-to-air temperature differences measured with infrared thermographers, extracted from multiple papers for a variety of geographical locations. Climate is denoted according to the Köppen–Geiger climate classification system (Peel et al., 2007)

| Species | $\Delta T_{canopy-air}$ | Site | Climate | Location | Reference |
|---|---|---|---|---|---|
| *Abies procera* | 5 | Urban | *Csb* | Oregon, USA | Kim et al. 2018 |
| *Acer platanoides* | 2 | City park | *Cfb* | Basel, Switzerland | Leuzinger et al. 2010 |
| *Acer rubrum* | 10.2 | Forest | *Dfb* | Massachusetts, USA | Aubrecht et al. 2016 |
| *Acer saccharinum* | 0.2 | City park | *Cfb* | Basel, Switzerland | Leuzinger et al. 2010 |
| *Aesculus hippocastanum* | -1.4 | City park | *Cfb* | Basel, Switzerland | Leuzinger et al. 2010 |
| *Betula papyrifera* | 6.7 | Forest | *Dfb* | Massachusetts, USA | Aubrecht et al. 2016 |
| *Carpinus betulus* | 10.6 | Closed-canopy forest | *Cfb* | Basel, Switzerland | Leuzinger and Körner 2007 |
| *Ceratonia siliqua* | 1.4 | Field | *Csa* | Beit Shemesh, Israel | Lapidot et al. 2019 |
| *Cupressus sempervirens* | 1.1 | Field | *Csa* | Beit Shemesh, Israel | Lapidot et al. 2019 |
| *Eucalyptus parramattensis* | -0.2 | Experimental setup | *Csa* / heatwave | NSW, Australia | Drake et al. 2018 |
| *Eucalyptus parramattensis* | 1.3 | Experimental setup | *Csa* | NSW, Australia | Drake et al. 2018 |
| *Fagus sylvatica* | 9.9 | Closed-canopy forest | *Cfb* | Basel, Switzerland | Leuzinger and Körner 2007 |
| *Gleditsia triacanthos* | -0.7 | City park | *Cfb* | Basel, Switzerland | Leuzinger et al. 2010 |
| *Larix decidua* | 4.5 | Closed-canopy forest | *Cfb* | Basel, Switzerland | Leuzinger and Körner 2007 |
| *Picea abies* | 8.8 | Closed-canopy forest | *Cfb* | Basel, Switzerland | Leuzinger and Körner 2007 |
| *Pinus halepensis* | -1.3 | Field | *Csa* | Beit Shemesh, Israel | Lapidot et al. 2019 |
| *Pinus ponderosa* | -2 | Closed-canopy forest | *Csb* | Oregon, USA | Kim et al. 2016 |
| *Pinus strobus* | 4.9 | Forest | *Dfb* | Massachusetts, USA | Aubrecht et al. 2016 |
| *Pinus sylvestris* | 1.3 | City park | *Cfb* | Basel, Switzerland | Leuzinger et al. 2010 |
| *Pinus sylvestris* | 5.4 | Closed-canopy forest | *Cfb* | Basel, Switzerland | Leuzinger and Körner 2007 |
| *Pinus virginiana* | 1.5 | Forest | *Cfa* | Virginia, USA | Yi et al. 2020 |
| *Pistacia lentiscus* | 3.1 | Field | *Csa* | Beit Shemesh, Israel | Lapidot et al. 2019 |
| *Platanus acerifolia* | 0.1 | City park | *Cfb* | Basel, Switzerland | Leuzinger et al. 2010 |
| *Prunus avium* | 6.4 | Closed-canopy forest | *Cfb* | Basel, Switzerland | Leuzinger and Körner 2007 |
| *Quercus alba* | 0.9 | Forest | *Cfa* | Virginia, USA | Yi et al. 2020 |
| *Quercus calliprinos* | 2.8 | Field | *Csa* | Beit Shemesh, Israel | Lapidot et al. 2019 |
| *Quercus falcata* | 1.6 | Forest | *Cfa* | Virginia, USA | Yi et al. 2020 |
| *Quercus petraea* | 7.2 | Closed-canopy forest | *Cfb* | Basel, Switzerland | Leuzinger and Körner 2007 |
| *Quercus rubra* | 10.5 | Forest | *Dfb* | Massachusetts, USA | Aubrecht et al. 2016 |
| *Robinia pseudoacacia* | 0.4 | City park | *Cfb* | Basel, Switzerland | Leuzinger et al. 2010 |
| *Tilia cordata* | 1.4 | City park | *Cfb* | Basel, Switzerland | Leuzinger et al. 2010 |
| *Tilia platyphyllos* | 2.6 | City park | *Cfb* | Basel, Switzerland | Leuzinger et al. 2010 |
| *Tilia platyphyllos* | 12 | Closed-canopy forest | *Cfb* | Basel, Switzerland | Leuzinger and Körner 2007 |
| *Tilia tomentosa* | 2.9 | City park | *Cfb* | Basel, Switzerland | Leuzinger et al. 2010 |
| Tropical trees | 1.9 | Rainforest | *Am* | Barro Colorado Island, Panama | Pau et al. 2018 |
